# Translating phenotypic prediction models from big to small anatomical MRI data using meta-matching

**DOI:** 10.1101/2023.12.31.573801

**Authors:** Naren Wulan, Lijun An, Chen Zhang, Ru Kong, Pansheng Chen, Danilo Bzdok, Simon B Eickhoff, Avram J Holmes, B. T. Thomas Yeo

**Affiliations:** Centre for Sleep & Cognition & Centre for Translational Magnetic Resonance Research, Yong Loo Lin School of Medicine, National University of Singapore; Department of Electrical and Computer Engineering, National University of Singapore, Singapore; N.1 Institute for Health & Institute for Digital Medicine, National University of Singapore, Singapore; Integrative Sciences and Engineering Programme (ISEP), National University of Singapore, Singapore; Department of Biomedical Engineering, McConnell Brain Imaging Centre (BIC), Montreal Neurological Institute (MNI), Faculty of Medicine, School of Computer Science, McGill University, Montreal QC, Canada; Mila – Quebec Artificial Intelligence Institute, Montreal, QC, Canada; Institute for Systems Neuroscience, Medical Faculty, Heinrich-Heine University Düsseldorf, Düsseldorf, Germany; Institute of Neuroscience and Medicine, Brain & Behavior (INM-7), Research Center Jülich, Jülich, Germany; Department of Psychiatry, Brain Health Institute, Rutgers University, Piscataway, NJ, USA; Martinos Center for Biomedical Imaging, Massachusetts General Hospital, Charlestown, MA, USA

**Keywords:** Structural MRI, transfer learning, meta-matching, phenotypic prediction

## Abstract

Individualized phenotypic prediction based on structural MRI is an important goal in neuroscience. Prediction performance increases with larger samples, but small-scale datasets with fewer than 200 participants are often unavoidable. We have previously proposed a “meta-matching” framework to translate models trained from large datasets to improve the prediction of new unseen phenotypes in small collection efforts. Meta-matching exploits correlations between phenotypes, yielding large improvement over classical machine learning when applied to prediction models using resting-state functional connectivity as input features. Here, we adapt the two best performing meta-matching variants (“meta-matching finetune” and “meta-matching stacking”) from our previous study to work with T1-weighted MRI data by changing the base neural network architecture to a 3D convolution neural network. We compare the two meta-matching variants with elastic net and classical transfer learning using the UK Biobank (N = 36,461), Human Connectome Project Young Adults (HCP-YA) dataset (N = 1,017) and HCP-Aging dataset (N = 656). We find that meta-matching outperforms elastic net and classical transfer learning by a large margin, both when translating models within the same dataset, as well as translating models across datasets with different MRI scanners, acquisition protocols and demographics. For example, when translating a UK Biobank model to 100 HCP-YA participants, meta-matching finetune yielded a 136% improvement in variance explained over transfer learning, with an average absolute gain of 2.6% (minimum = -0.9%, maximum = 17.6%) across 35 phenotypes. Overall, our results highlight the versatility of the meta-matching framework.

## 1. Introduction

An important goal in systems neuroscience is to understand how variation in brain structure relates to individual differences in behavior (Genon et al., 2022). Structural T1-weighted magnetic resonance imaging (MRI) is a non-invasive technique for examining the anatomy of the human brain, providing high contrast between gray and white matter (Gifford et al., 2020). Structural MRI is widely used to predict behavioral traits, clinical symptoms and diagnostic categories in both healthy individuals and individuals with neuropsychiatric disorders (Sabuncu et al., 2015; Arbabshirani et al., 2017; Bhagwat et al., 2019; Cohen et al., 2021; Ooi et al., 2022). However, most prediction studies use datasets with fewer than a few hundred participants, leading to low reproducibility and inflated performance (Arbabshirani et al., 2017; Bzdok et al., 2018; Masouleh et al., 2019; Poldrack et al., 2020; Marek et al., 2022). Studies have shown that prediction performance increases with larger sample sizes (Chu et al., 2012; Cui & Gong, 2018; He et al., 2020; Schulz et al., 2020), but for investigations of certain clinical populations or focused neuroscience inquiries, small-scale datasets remain unavoidable. Here, to address this fundamental issue, we seek to establish a framework to translate prediction models from large-scale datasets to predict new non-brain-imaging phenotypes in small-scale datasets based on anatomical T1-weighted images.

More specifically, given a large-scale anatomical MRI dataset (N > 10,000) with multiple phenotypes, we seek to translate models trained from the large dataset to new unseen phenotypes in a small independent dataset (N ≤ 200). In machine learning, this problem is often referred to as meta-learning, lifelong learning, learning-to-learn or few-shot learning (Fei-Fei et al., 2006; Andrychowicz et al., 2016; Finn et al., 2017; Ravi & Larochelle, 2016; Vanschoren, 2019), and is closely related to transfer learning (Weiss et al., 2016; Hospedales et al., 2021). Broadly speaking, meta-learning and transfer learning methods usually train a model on abundant data on a related problem, called the source dataset, and seek to translate knowledge learned from the large-scale dataset to the small dataset, called the target dataset. During the translation, a subset of the target dataset is typically used to adapt the pre-trained model to the new sample. One distinction between meta-learning and transfer learning is that in transfer learning, the prediction problem in the target dataset can be same (Ghafoorian et al., 2017; Aderghal et al., 2018; Wee et al., 2019) or different (Dawud et al., 2019; Talo et al., 2019; Mehmood et al., 2021) from the source dataset. On the other hand, meta-learning always involves the translation of the prediction model to perform a *new* prediction problem in the target dataset – providing the imaging neuroscience community with a versatile modeling framework that, once established, can be applied to a diversity of research goals.

In our previous study (He et al., 2022), we developed a simple “meta-matching” approach to translate prediction models from large datasets to improve the prediction of new phenotypes in small datasets. Meta-matching is grounded in the observation that many phenotypes are correlated, as demonstrated by previous studies identifying a small number of factors linking brain imaging data to various non-brain-imaging traits like cognition, mental health, demographics, and other health attributes (Smith et al., 2015; Miller et al., 2016; Xia et al., 2018; Kebets et al., 2019). As a result, a phenotype X in a smaller-scale study is likely correlated, sharing a latent relationship, with a phenotype Y present in a larger population dataset. Therefore, a model trained to predict phenotype Y from the larger dataset might be predisposed to features useful for predicting phenotype X. Consequently, the predictive model of Y can be more effectively translated to predict phenotype X in the smaller study. As a demonstration of meta-matching (He et al., 2022), we trained a simple fully-connected feedforward neural network to predict 67 non-brain-imaging phenotypes from resting-state functional connectivity (RSFC) in the UK Biobank. The neural network was then translated using meta-matching to predict non-brain-imaging phenotypes in the Human Connectome Project Young Adult (HCP-YA) dataset, yielding large improvements in prediction accuracies over classical kernel ridge regression (without meta-learning or transfer learning).

In the current study, we investigated whether the two best performing meta-matching variants (“meta-matching finetune” and “meta-matching stacking”) from our previous study (He et al., 2022) can be adapted to work with T1 MRI data. More specifically, given the different modalities (RSFC versus T1), the base neural network architecture was changed from a fully-connected feedforward neural network to the simple fully convolutional network (SFCN; Peng et al., 2021). The SFCN was chosen because of its simplicity and top performance in the Predictive Analysis Challenge 2019 of brain age prediction (Peng et al., 2021). We compared the two meta-matching variants with classical elastic net and classical transfer learning using the UK Biobank (Sudlow et al., 2015; Miller et al., 2016), Human Connectome Project Young Adults (HCP-YA) dataset (Van Essen et al., 2013) and HCP-Aging dataset (Harms et al., 2018; Bookheimer et al., 2019).

It is worth mentioning that it is not obvious that meta-matching will confer great benefits in anatomical MRI, compared with RSFC (He et al., 2022). The reason is that RSFC-based prediction typically utilizes high dimensional features derived from N × N RSFC matrices, where N is the number of brain parcels (or independent component analysis components). On the other hand, T1-based prediction can utilize low dimensional N × 1 volumetric and/or thickness measures. Therefore, classical machine learning techniques (e.g., elastic net) might work really well in the small sample regime (≤ 200 participants). Nevertheless, we found that meta-matching significantly outperformed classical elastic net and transfer learning, highlighting the versatility of the meta-matching framework.

## 2. Methods

### 2.1. Datasets and preprocessing

In this section, we describe the datasets and preprocessing used in the current study. Table 1 summarizes the demographics and acquisition parameters of the three datasets we considered. We will evaluate meta-matching based on prediction accuracy when translating prediction models within the same dataset (UK Biobank), as well as across datasets, i.e., from UK Biobank to HCP-YA and HCP-Aging datasets. The very different age ranges between HCP-YA and UK Biobank served as a strong test of the generalizability of meta-matching. All data collection and analysis procedures were approved by the respective institutional review boards (IRBs), including the National University of Singapore IRB for the analysis presented in this paper.

**Table 1.** Summary of demographics and acquisition parameters of the three datasets used in the current study.

|  | Age | Sex (M/F) | Scanner(s) | Resolution |
| --- | --- | --- | --- | --- |
| UK Biobank | 45-82 | 53%/47% | Siemens Skyra<br>3T scanner | 1mm |
| HCP-YA | 22-35 | 47%/53% | Customized<br>Skyra 3T scanner | 0.7mm |
| HCP-Aging | 36-100 | 44%/56% | Siemens Prisma<br>3T scanner | 0.8mm |

#### 2.1.1. UK Biobank

The UK Biobank (UKBB) dataset is a large-scale epidemiology study of over 500,000 adults from the United Kingdom (Alfaro-Almagro et al., 2018). The volunteers were recruited between 2006 and 2010 from 22 centers across the UK. Participants were asked to answer a variety of questionnaires about different aspects of health and lifestyle. In addition, a range of physiological measurements was also collected. We considered the same set of 36,848 participants and 67 non-brain-imaging phenotypes (referred to as phenotypes henceforth; Table S1) from our previous study (He et al., 2022).

As part of the UK Biobank pipeline (Alfaro-Almagro et al., 2018), FreeSurfer recon-all was used to derive thickness and volume measures with the Desikan-Killiany-Tourville (DKT40) cortical atlas (Klein et al., 2012) and subcortical segmentation (Fischl et al., 2002). We considered the subset of regions present in most participants, yielding 164 morphometric measures, comprising intracranial volume (ICV) and thickness measures of 62 cortical regions, as well as volumes of 62 cortical regions and 39 subcortical regions (Tables S4 and S5). After excluding participants who have dropped out since our previous study (He et al., 2022) and excluding participants without all 164 morphometric measures, we ended up with 36,461 participants. As a baseline, these 164 measures will be utilized by the elastic net algorithm for phenotypic prediction (see Section 2.3).

Furthermore, we used FMRIB’s Linear Image Registration Tool (FLIRT) to transform the bias-field-corrected version of the brain-extracted T1 (from the UK Biobank provided preprocessing outputs) to MNI152 standard-space T1 template with 1 mm resolution (Jenkinson et al., 2001; Jenkinson et al., 2002). Each T1 image was cropped to dimensions 160 x 192 x 160, and then divided by the mean value within each image following Peng et al. (2021). The normalized T1 images will be used by a convolutional neural network for phenotypic prediction (Section 2.2).

#### 2.1.2. HCP Young Adult (HCP-YA) dataset

We utilized the Human Connectome Project Young Adult (HCP-YA) dataset (Van Essen et al., 2013), which included healthy young adults. We considered 1,019 participants and 35 non-brain-imaging phenotypes, consistent with our previous study (He et al., 2022). The phenotypes are found in Table S2.

FreeSurfer recon-all from the HCP pipeline was used to derive thickness and volume measures with the DKT40 cortical atlas (Klein et al., 2012) and ASEG subcortical segmentation (Fischl et al., 2002). We considered the subset of regions present in most participants, yielding 166 morphometric measures, comprising intracranial volume (ICV) and thickness measures of 62 cortical regions, as well as volumes of 62 cortical regions and 41 subcortical regions (Tables S4 and S5). We note that the difference in the number of morphometric measures between UK Biobank and HCP-YA datasets (164 vs 166) arose because the 5th-Ventricle and non-WM-hypointensities were missing in most participants from the UK Biobank dataset. As a baseline, these 166 measures will be utilized by the elastic net algorithm for phenotypic prediction (see Section 2.2).

Moreover, we considered T1 images of 0.7 mm resolution which had been transformed to MNI152 space by FLIRT from the HCP PreFreesurfer pipeline. We noted the files of two participants were missing in the HCP filesystem, so we ended up with 1,017 participants. Each T1 image was downsampled to 1mm, cropped to dimensions 160 x 192 x 160, and then divided by the mean value within each image following Peng et al. (2021). The processed T1 images will be used by a convolutional neural network for phenotypic prediction (Section 2.2).

#### 2.1.3. HCP-Aging dataset

Besides the HCP-YA dataset, we also used the Human Connectome Project Aging (HCP-Aging) dataset (Harms et al., 2018; Bookheimer et al., 2019) consisting of healthy participants. We manually selected commonly used non-brain-imaging phenotypic measures across cognition, emotion, motor, sensor, and life experience, resulting in 45 phenotypes (Table S3). By only considering participants with at least 90% of the phenotypes, we ended up with 656 participants (out of 725 participants). Similar to the HCP-YA dataset, we used the same 166 morphometric measures generated by the FreeSurfer recon-all procedure from the HCP pipeline. Moreover, we considered T1 images of 0.8 mm resolution, which had been transformed to MNI152 space by FLIRT from the HCP PreFreesurfer pipeline. Each T1 image was downsampled to 1mm, cropped to dimensions 160 x 192 x 160, and then divided by the mean value within each image following Peng et al. (2021). The processed T1 images will be used by a convolutional neural network for phenotypic prediction (Section 2.2).

### 2.2. Data split for different analyses

We performed two sets of analyses. First, we benchmarked meta-matching within the UK Biobank. Second, we translated predictive models from the UK Biobank to the HCP-YA and HCP-Aging datasets.

#### 2.2.1. Data split within UK Biobank

For the UK Biobank analysis, we considered 36,461 participants with T1 structural MRI and 67 phenotypes. As illustrated in Figure 1, we randomly split the data into a meta-training set comprising 26,573 participants with 33 phenotypes, as well as a meta-test set comprising 9,888 participants with 34 phenotypes. There was no overlap between the participants and phenotypes across the meta-training set and meta-test set.

**Figure 1.**
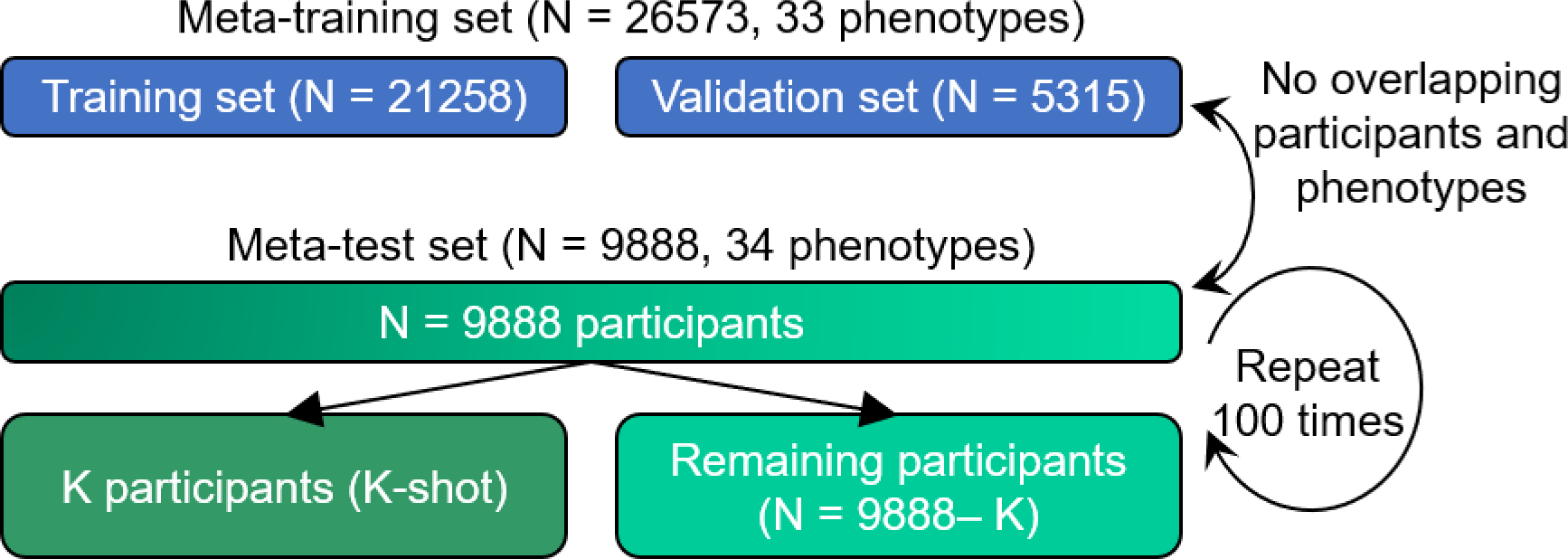
Data split scheme for within-UK Biobank analysis. The UK Biobank dataset was divided into a meta-training set comprising 26,573 participants and 33 phenotypes, as well as a meta-test set comprising 9,888 participants and 34 other phenotypes. There was no participant or phenotype overlap between meta-training and meta-test sets. The meta-test set was, in turn, split into K participants (K = 10, 20, 50, 100 and 200) and remaining 9,888 − K participants. The group of K participants mimicked studies with traditionally common sample sizes. Various trained models from the meta-training set were translated to the meta-test set using the K participants. The models were then evaluated using the remaining N – K participants. This random split was repeated 100 times for robustness.

We further randomly split the meta-training set into a training set with 21,258 participants (80% of 26,573 participants) and a validation set with 5,315 participants (20% of 26,573 participants). The validation set was used for tuning hyperparameters of the predictive models.

For the meta-test set, we randomly split 9,888 participants into K participants (K-shot) and 9,888− K participants, where K had a value of 10, 20, 50, 100, and 200. The group of K participants mimicked traditional small-N studies. Various trained models from the meta-training set were translated to the meta-test set using the K participants. The models were then evaluated using the remaining N – K participants. Each random K-shot split was repeated 100 times to ensure stability.

#### 2.2.2. Data split scheme for cross-dataset analyses

To translate predictive models from the UK Biobank to other datasets, we considered HCP-YA and HCP-Aging datasets. As illustrated in Figure 2, the meta-training set comprised all 36,461 participants with all 67 phenotypes from the UK Biobank dataset. The first meta-test set consisted of 1,017 participants with 35 phenotypes from the HCP-YA dataset. The second meta-test set consisted of 656 participants with 45 phenotypes from the HCP-Aging dataset. There was no overlap between the participants and phenotypes across the meta-training and meta-test sets because they were from totally different datasets. For the meta-training set, we further randomly split it into a training set with 29,169 participants (80% of 36,461 participants) and a validation set with 7,292 participants (20% of 36,461 participants). The validation set was used for tuning hyperparameters of the predictive models.

**Figure 2.**
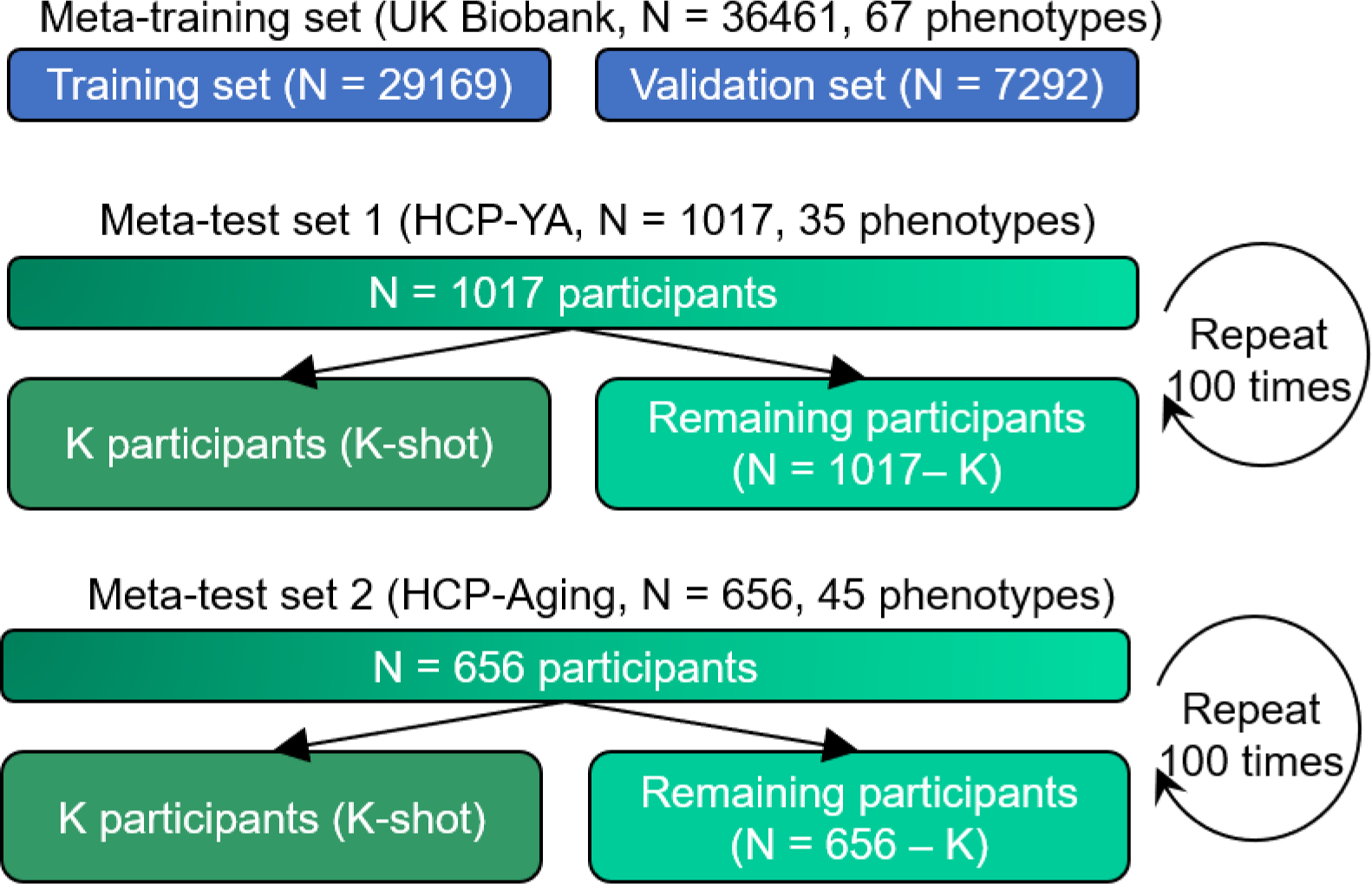
Data split scheme for cross-dataset analysis. The meta-training set comprised 36,461 UK Biobank participants and 67 phenotypes. The first meta-test set comprised 1,017 HCP-YA participants and 35 phenotypes. The second meta-test set comprised 656 HCP- Aging participants and 45 phenotypes. Each meta-test was, in turn, split into K participants (K = 10, 20, 50, 100 and 200) and remaining participants. The group of K participants mimicked studies with traditionally common sample sizes. Various trained models from the meta-training set were translated to the meta-test set using the K participants. The models were then evaluated using the remaining N – K participants. This random split was repeated 100 times for robustness.

For the HCP-YA dataset, we randomly split 1,017 participants into K participants (K-shot) and 1,017− K participants, where K had a value of 10, 20, 50, 100, and 200. Various trained models from the meta-training set were translated to the meta-test set using the K participants. The models were then evaluated using the remaining N – K participants. Each random K-shot split was repeated 100 times to ensure stability. The same procedure was applied to the HCP-Aging dataset.

### 2.3. Predictive models

Figure 3 provides an overview of the different approaches we will compare. Across all approaches, we z-normalize each non-imaging phenotype to have zero mean and unit variance across participants. More specifically, in the case of the meta-training set, the mean and standard deviation were computed using all the participants to apply the z-normalization. In the case of the meta-test set, for each phenotype, the mean and standard deviation were computed from the K participants and subsequently carried over to the full meta-test set comprising the K participants and the remaining N – K test participants.

**Figure 3.**
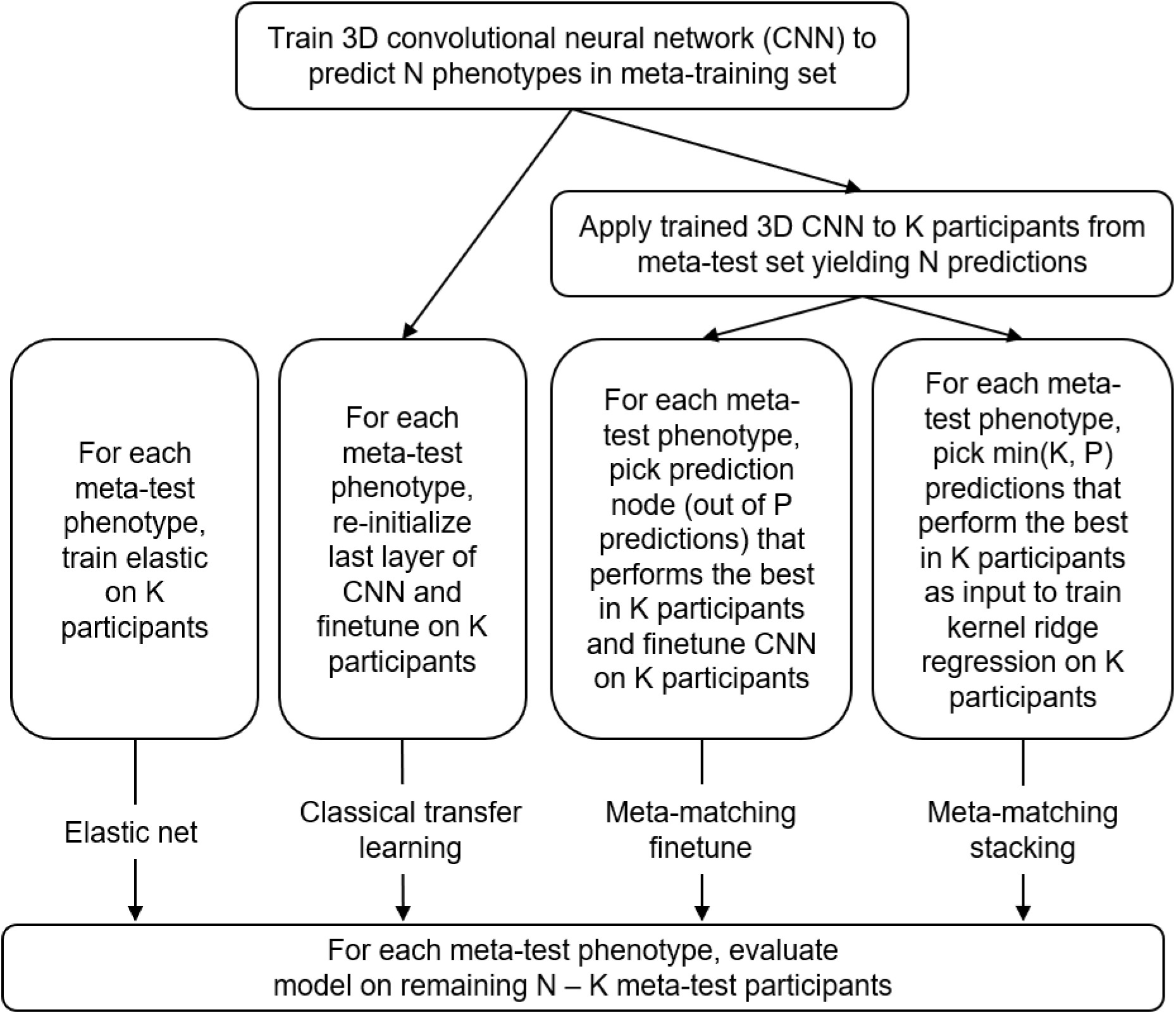
Overview of different approaches. We considered two baselines: elastic net and classical transfer learning. We proposed two meta-matching variants: meta-matching finetune and meta-matching stacking.

Following our previous study (He et al., 2022), statistical difference between algorithms was evaluated using a bootstrapping approach (more details in Supplementary Methods S2). More specifically, we will compare both meta-matching variants (Sections 2.3.3 and 2.3.4) with the two baselines (Sections 2.3.1 and 2.3.2). Multiple comparisons were corrected using a false discovery rate (FDR) of q < 0.05. FDR was applied to all K-shots, across all comparisons and both evaluation metrics (Pearson’s correlation and COD).

#### 2.3.1. Baseline 1: Elastic net

As a baseline, we used thickness and volumetric measures as input features to predict individuals’ phenotypes using elastic net (Figure 3). Elastic net is a linear regression model with an L1 lasso and L2 ridge regularization terms (Zou et al., 2005). Here, we chose elastic net as a baseline because previous studies have suggested that elastic net yielded strong prediction performance in phenotypic prediction for brain MRI data (Pervaiz et al., 2020; Peng et al., 2021; Ooi et al., 2022).

Given K participants from the meta-test set, the morphometric (volumetric and thickness) measures were z-normalized based on the mean and standard deviation computed from the K participants. We note that the morphometric measures of the N – K participants were also z-normalized using the mean and standard deviation computed from the K participants. The z-normalized morphometric measures were used as input to train the elastic net model on the K participants.

More specifically, we performed five-fold cross-validation on the K participants with different combinations of the hyperparameters *λ*_1_ and *λ*_2_(which controlled the strength of the L1 and L2 regularizations). We used coefficient of determination (COD) to evaluate prediction performance to choose the best hyperparameters for *λ*_1_ and *λ*_2_ across the 5-fold cross-validation.

The best hyperparameters *λ*_1_ and *λ*_2_were then used to train the elastic net model using all K participants. The trained elastic net model was then applied to the remaining N – K test participants in the meta-test set. Pearson’s correlation and the COD were used to evaluate prediction performance. This procedure was repeated for each of the 100 random splits.

#### 2.3.2. Baseline 2: Classical transfer learning

To perform classical transfer learning, we first trained a simple fully convolutional network (SFCN) introduced by Peng et al., (2021) in the meta-training set to jointly predict all the available meta-training phenotypes.

The input to the SFCN is the mean-normalized T1 image affine transformed to MNI152 standard space (Section 2.1). The SFCN’s convolutional neural network (CNN) architecture was based on VGG (Simonyan et al., 2014) and used a fully convolutional structure (Long et al., 2015). We chose the SFCN given its simplicity and top performance in the Predictive Analysis Challenge 2019 of brain age prediction (Peng et al., 2021). In the original study (Peng et al., 2021), the last layer comprised 40 nodes that represented the predicted probability of the age interval that a participant’s age falls into. Here, we modified the last layer to predict P phenotypes simultaneously. P is equal to 33 in the within-UK Biobank analysis (Figure 1) and P is equal to 67 in the cross-dataset analysis (Figure 2).

Figure 4 shows the overall network architecture. The 3D CNN consisted of several convolutional blocks for feature extraction. Each feature extraction block (except the last block) consisted of a 3D convolutional layer, a batch normalization layer, a max pooling layer, and a ReLU activation layer. The last block was similar to the previous blocks but without the max pooling layer. The feature maps from the last block were fed into an average pooling layer (green in Figure 4).

**Figure 4.**
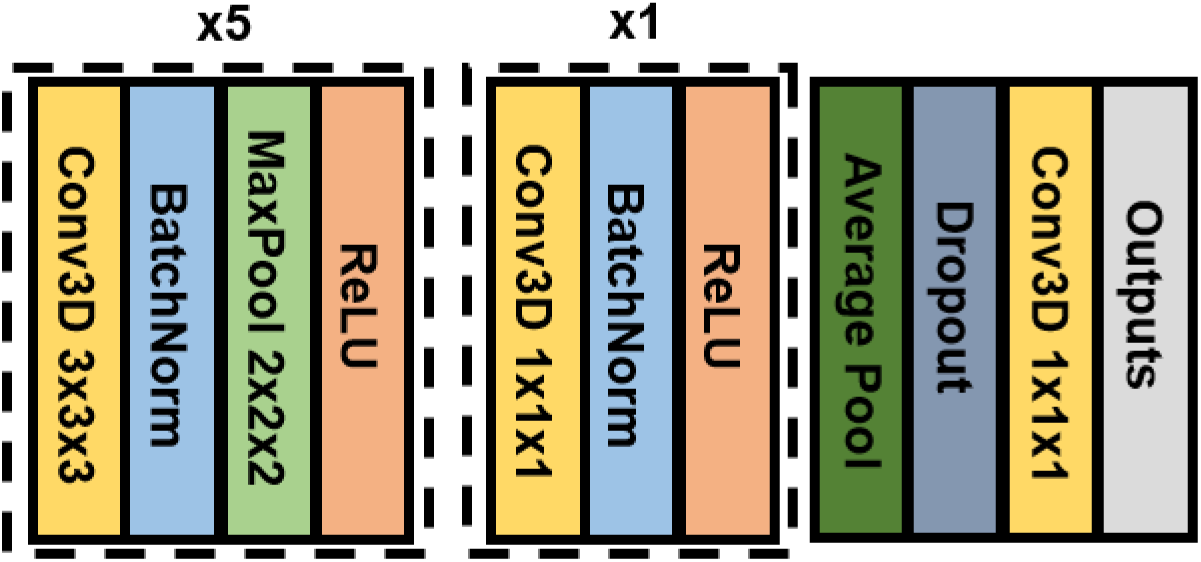
Network architecture of the Simple Fully Convolutional Neural Network (SFCN) model (Peng et al., 2021) adapted to the current study. In the original study (Peng et al., 2021), the last layer comprised 40 nodes that represented the predicted probability of the age interval that a participant’s age falls into. Here, we modified the last layer to predict P phenotypes simultaneously.

Since the elastic net utilized ICV as one of the features, while affine registration of T1 to MNI152 space removed this information, for the comparison to be comparable, we concatenated z-normalized ICV with the outputs of the average pooling layer. More specifically, for both meta-training and meta-test sets, ICV of each participant was z-normalized using the mean and standard deviation computed from the participants of the training set within the meta-training set. The concatenated features were then fed into a dropout layer and then went through a 3D convolution layer with 1×1×1 kernel size to produce the final outputs.

The hyperparameters of the CNN were empirically determined based on the meta-training set from the within UK Biobank analysis (Figure 1A). Both within-UK Biobank and cross-dataset analyses used the same set of hyperparameters. More details about model architecture and hyperparameters (for example, the number of blocks, number of channels per block, and kernel size per block) can be found in Supplementary Methods S1.

The same transfer learning procedure was used for both within-UK Biobank analysis and cross-dataset analysis (Figure 3). The only difference is that the within-UK Biobank analysis used a CNN model trained on 26,573 participants and 33 phenotypes, while the cross-dataset analysis used a CNN model trained on 36,461 participants and 67 phenotypes.

To perform transfer learning, we first replaced the last layer of the 3D CNN model (trained on the meta-training set) with a new convolutional layer with 1×1×1 kernel size and one output node. The new convolutional layer was initialized with random weights. For each meta-test phenotype, the last two layers of the CNN model were then finetuned on K participants in the meta-test set, while the weights of the remaining layers were frozen.

The optimal learning rate was determined using grid search and five-fold cross-validation on the K participants. The optimal learning rate was then used to perform fine-tune a final model using all K participants. For both the five-fold cross validation and the final round of fine-tuning, the maximum fine-tuning epochs was set to be 50 with 80% of K participants used for training and 20% used to evaluate validation loss for early stopping, to reduce the possibility of overfitting. This final trained model was evaluated in the remaining N – K participants in the meta-test set. Pearson’s correlation and COD were used to evaluate the prediction performance. This procedure was repeated for each of the 100 random splits.

#### 2.3.3. Meta-matching finetune

As an alternative to transfer learning, we considered the “meta-matching finetune” approach (Figure 3) introduced in our previous study (He et al., 2022). To explain the meta-matching finetune procedure in the current study, we will focus on the experimental setup for the within-UK Biobank analysis.

Recall from Section 2.3.2 that we have trained a 3D CNN model to predict 33 phenotypes in the meta-training set from the UK Biobank. Given K participants from the meta-test set, we applied the CNN yielding 33 predictions. For each meta-test phenotype (out of 34 phenotypes), we calculated the accuracy (COD) with each of the 33 predictions for the K participants. The output node of the CNN model with the best COD was chosen, while the remaining 32 nodes were removed. The last two layers of the CNN model were finetuned using the K participants, while the weights of the remaining layers were frozen.

Therefore, the difference between meta-matching finetune and classical transfer learning (Section 2.3.2) is the initialization of the last layer. Classical transfer learning randomly initialized the last layer, while meta-matching finetune initialized the last layer by choosing the “closest” phenotypic prediction model from the meta-training set.

The optimal learning rate for finetuning was determined using grid search and five-fold cross-validation on the K participants. The optimal learning rate was then used to perform fine-tune a final model using all K participants. For both the five-fold cross validation and the final round of fine-tuning, the maximum fine-tuning epochs was set to be 50 with 80% of K participants used for training and 20% used to evaluate validation loss for early stopping, to reduce the possibility of overfitting.

This final trained model was evaluated in the remaining N – K participants in the meta-test set. Pearson’s correlation and COD were used to evaluate the prediction performance. This procedure was repeated for each of the 100 random splits. The same procedure was used for both within-UK Biobank analysis and cross-dataset analysis (Figure 3). The only difference was that the within-UK Biobank analysis used a CNN model trained on 26,573 participants and 33 phenotypes, while the cross-dataset analysis used a CNN model trained on 36,461 participants and 67 phenotypes.

#### 2.3.4. Meta-matching stacking

We also considered the meta-matching stacking approach (Figure 3) introduced in our previous study (He et al., 2022). To explain the meta-matching stacking procedure in the current study, we will again focus on the experimental setup for the within-UK Biobank analysis.

Recall from Section 2.3.2 that we have trained a 3D CNN model to predict 33 phenotypes in the meta-training set from the UK Biobank. Given K participants from the meta-test set, we applied the CNN yielding 33 predictions. For each meta-test phenotype (out of 34 phenotypes), we calculated the accuracy (COD) with each of the 33 predictions for the K participants, and selected the top M predictions. The value of M was set to be the minimum of K and 33 to reduce overfitting. For example, when K = 20, then M was set to be 20. When K = 50, then M was set to be 33.

A stacking procedure was then performed (Wolpert, 1992; Breiman, 1996), in which a kernel ridge regression (KRR) model was trained on K participants using the M predictions as input to predict the meta-test phenotype. Similar to our previous study (He et al., 2022), we used the correlation kernel. The hyperparameter λ was tuned using grid search and five-fold cross-validation on the K participants. The optimal λ was then used to train a final KRR model using all K participants.

The trained KRR model was then applied to the remaining N – K participants in the meta-test set. Pearson’s correlation and the COD were used to evaluate the prediction performance. This procedure was repeated for each of the 100 random splits.

### 2.4. Deep neural network implementation

The deep neural network was implemented using PyTorch (Paszke et al., 2017) and computed on NVIDIA RTX 3090 GPUs with CUDA 11.0. More details can be found in Supplementary Methods S1.

### 2.5. Model interpretation

To illustrate how meta-matching models can be interpreted, similar to our previous study (He et al., 2022), we utilized Haufe transform (Haufe et al., 2014) to interpret the meta-matching stacking prediction of Rey Auditory Verbal Learning Test (RAVLT) score and Montreal Cognitive Assessment (MOCA) in the HCP-Aging dataset for K = 100 participants.

For a predictive model with T1 structural MRI as input and phenotype as output, Haufe transform produces a feature importance value for each voxel. A positive (or negative) predictive feature value implied that higher T1 intensity was related to predicting greater (or lower) phenotypic score.

More specifically, for each phenotype, Haufe transform was calculated as the covariance between the phenotype’s prediction based on the meta-matching stacking model and the intensity value of each T1 voxel (across the 100 participants), yielding a 3D volume. The 3D volumes were averaged across the 100 random sampling of 100 participants, and were then visualized in MNI152 space.

### 2.6. Data and code availability

The code can be found here (https://github.com/ThomasYeoLab/CBIG/XXX). Two co-authors (Lijun An and Chen Zhang) reviewed the code before merging it into the GitHub repository to reduce the chance of coding errors. The trained models for meta-matching are also publicly available (https://github.com/ThomasYeoLab/CBIG/tree/master/stable_projects/predict_phenotypes/N <u>WULAN2024_MMS</u>).

This study used publicly available data from the UK Biobank (https://www.ukbiobank.ac.uk/), as well as the HCP-YA and HCP-Aging datasets (https://www.humanconnectome.org/). Data can be accessed via data use agreements.

## 3. Results

3.1. Meta-matching outperforms elastic net and transfer learning within the UK Biobank Four approaches (elastic net, classical transfer learning, meta-matching finetune, and meta-matching stacking) were applied to the UK Biobank to predict 34 meta-test phenotypes. The models were trained or adapted based on K participants and then evaluated on the remaining 9,888 – K participants (Figure 1).

Figures 5A and 6A show the Pearson’s correlation and COD respectively, averaged across all 34 meta-test phenotypes. Each boxplot represents 100 random samplings of K participants. Figures 5B and 6B show the outcomes of the statistical tests obtained by a bootstrapping procedure (Supplementary Methods S2). The actual p values are reported in Table S6. Colors indicate effect sizes of differences (Cohen’s D) between approaches.

**Figure 5.**
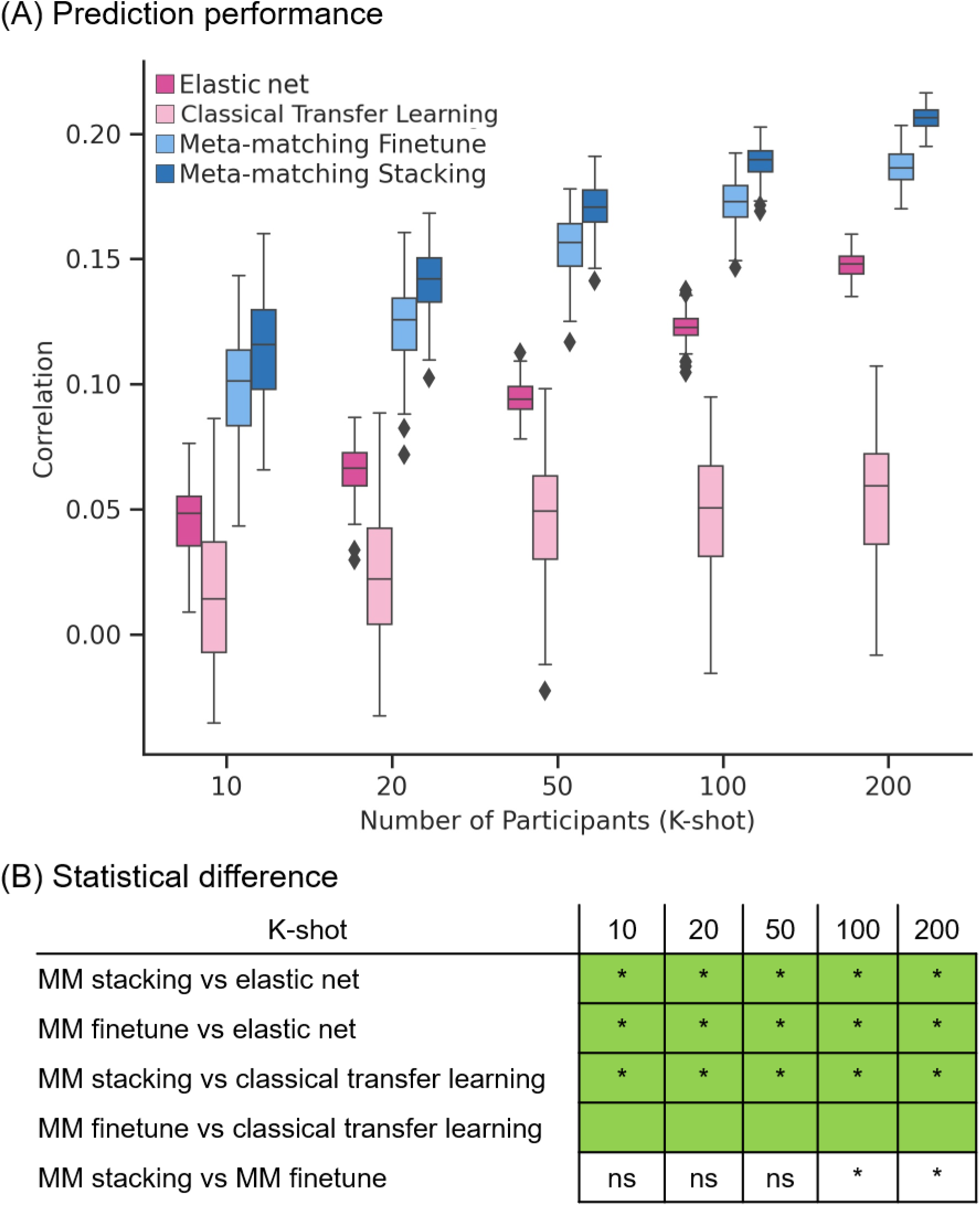
Meta-matching compared favorably with elastic net and direct transfer learning in terms of Pearson’s correlation within the UK Biobank. (A) Phenotypic prediction performance (Pearson’s correlation) averaged across 34 meta-test phenotypes in the UK Biobank. X-axis is the number of participants in the meta-test set of the UK Biobank used to train an elastic net or adapt the pretrained model from the meta-training set of the UK Biobank. Each boxplot shows the distribution of performance over 100 repetitions of sampling K participants. (B) Statistical difference between the prediction performance (Pearson’s correlation) of baseline methods and meta-matching algorithms. P values were calculated based on a two-sided bootstrapping test. ‘*’ indicates statistical significance after multiple comparisons correction (FDR q < 0.05). ‘ns’ indicates statistical test did not survive FDR correction. We note that there was no statistical test between meta-matching finetune and classical transfer learning because the bootstrapping procedure was too expensive for the two methods. Colors indicate effect sizes of differences (Cohen’s D) between approaches. Light green color indicates effect size **≥** 0.8. Dark green color indicates 0 ≤ effect size < 0.8. Dark pink indicates ™0.8 < effect size < 0. Light pink color indicates effect size ≤ ™0.8. There is no color for the comparison between meta-matching finetune and stacking since they are both our proposed methods.

**Figure 6.**
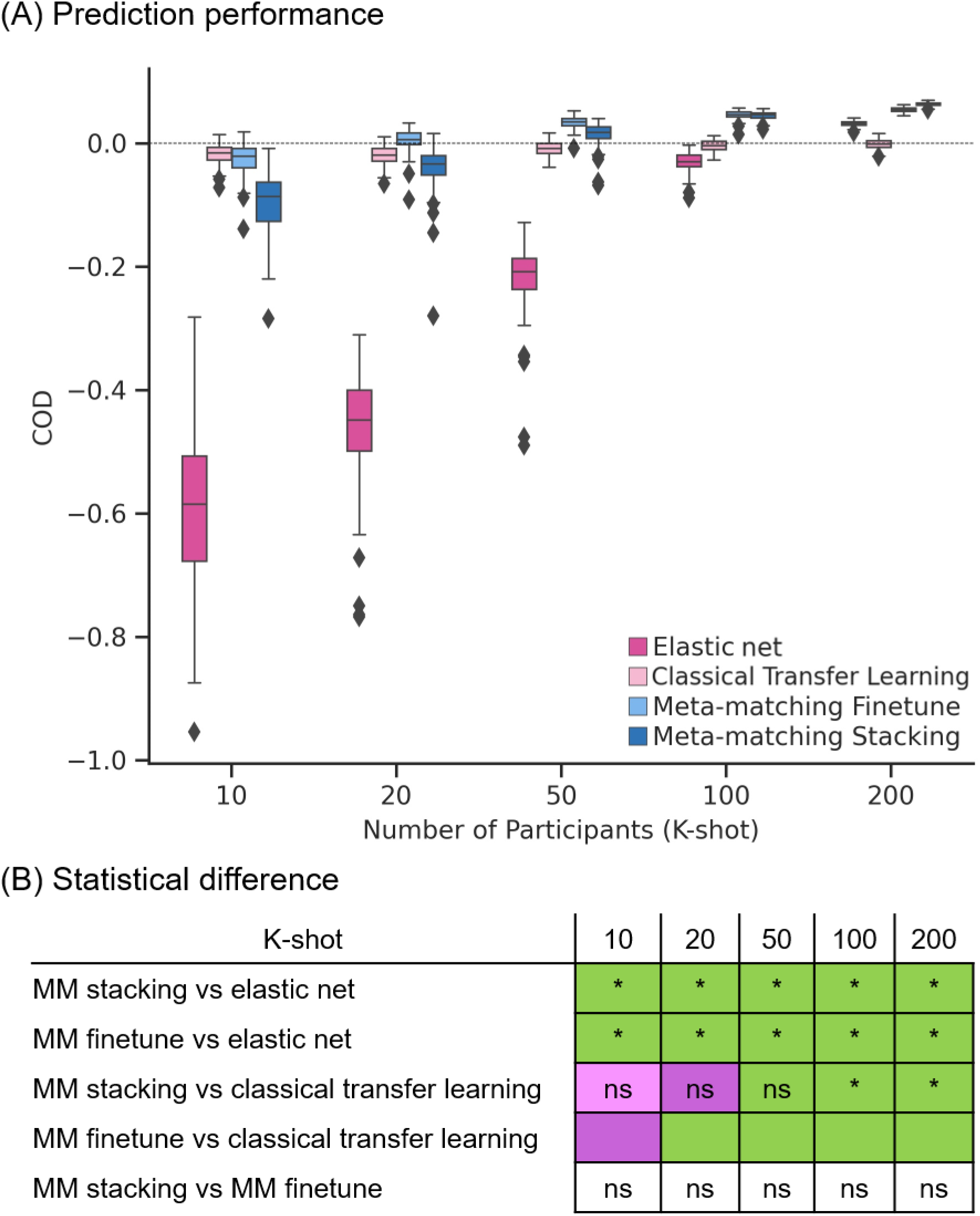
Meta-matching compared favorably with elastic net and direct transfer learning in terms of coefficient of determination (COD) within the UK Biobank. (A) Phenotypic prediction performance (COD) averaged across 34 meta-test phenotypes in the UK Biobank. X-axis is the number of participants in the meta-test set of the UK Biobank used to train an elastic net or adapt the pretrained model from the meta-training set of the UK Biobank. Each boxplot shows the distribution of performance over 100 repetitions of sampling K participants. (B) Statistical difference between the prediction performance (COD) of baseline methods and meta-matching algorithms. P values were calculated based on a two-sided bootstrapping test. ‘*’ indicates statistical significance after multiple comparisons correction (FDR q < 0.05). ‘ns’ indicates statistical test did not survive FDR correction. We note that there was no statistical test between meta-matching finetune and classical transfer learning because the bootstrapping procedure was too expensive for the two methods. Colors indicate effect sizes of differences (Cohen’s D) between approaches. Light green color indicates effect size ≥ 0.8. Dark green color indicates 0 ≤ effect size < 0.8. Dark pink indicates ™0.8 < effect size < 0. Light pink color indicates effect size ≤ ™0.8. There is no color for the comparison between meta-matching finetune and stacking since they are both our proposed methods.

In the case of Pearson’s correlation (Figure 5), both meta-matching finetune and meta-matching stacking greatly outperformed elastic net and classical transfer learning for all values of K. Meta-matching stacking was statistically better than meta-matching finetune for K **≥** 100.

In the case of COD (Figure 6), both meta-matching finetune and meta-matching stacking greatly outperformed elastic net for all values of K. For K ≤ 20, classical transfer learning was numerically better but not statistically better than meta-matching stacking. From K **≥** 50, meta-matching stacking was numerically better than transfer learning with statistical significance from K = 100 onwards.

On the other hand, for K = 10, classical transfer learning was numerically better than meta-matching finetune, while meta-matching finetune was better than classical transfer learning for remaining other values of K with large effect sizes (light green in Figure 6B). We note that there was no statistical test between meta-matching finetune and classical transfer learning because of the huge computational cost of the two approaches, so no bootstrapping was performed for either approach.

Another relevant point is that COD for all approaches were negative for K = 10. COD was positive for meta-matching finetune for K = 20 onwards, and positive for meta-matching stacking for K = 50 onwards. This suggests that absolute prediction accuracy (i.e., COD) is difficult even with meta-learning or transfer learning, when the sample size is very small.

Overall, meta-matching was better than elastic net for all values of K for both evaluation metrics (Pearson’s correlation and COD). On the other hand, meta-matching compared favorably with respect to transfer learning for all values of K for Pearson’s correlation and for larger values of K for COD.

### 3.2. Meta-matching outperforms baselines in the HCP-YA dataset

The previous experiment results (Figures 5 and 6) suggest that meta-matching can perform well when transferring within the same dataset (UK Biobank). We now evaluate the generalizability of meta-matching across datasets, using the HCP-YA and HCP-Aging datasets (Figure 2) in the following section and next section respectively.

Figures 7A and 8A show the Pearson’s correlation and COD respectively, averaged across all 35 meta-test phenotypes in the HCP-YA dataset. Each boxplot represents 100 random samplings of K participants. Figures 7B and 8B show the outcomes of the statistical tests obtained by a bootstrapping procedure (Supplementary Methods S2). The actual p values are reported in Table S7. Colors indicate effect sizes of differences (Cohen’s D) between approaches.

**Figure 7.**
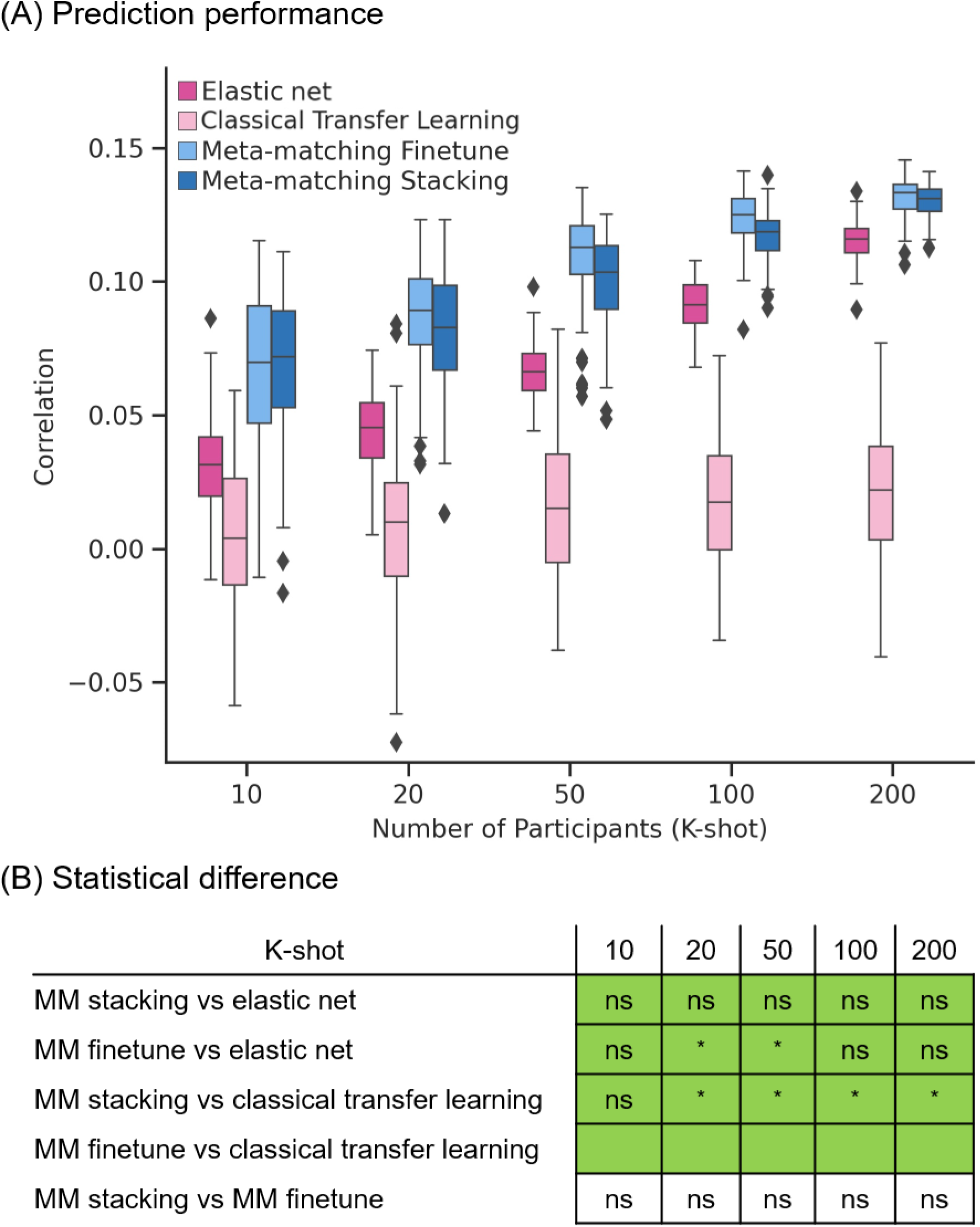
Meta-matching compared favorably with elastic net and direct transfer learning in terms of Pearson’s correlation when translating models from UK Biobank to HCP-YA dataset. (A) Phenotypic prediction performance (Pearson’s correlation) averaged across 35 meta-test phenotypes in the HCP-YA dataset. X-axis is the number of participants from the HCP-YA dataset used to train an elastic net or adapt the pretrained model from the meta-training set. Each boxplot shows the distribution of performance over 100 repetitions of sampling K participants. (B) Statistical difference between the prediction performance (Pearson’s correlation) of baseline methods and meta-matching algorithms. P values were calculated based on a two-sided bootstrapping test. ‘*’ indicates statistical significance after multiple comparisons correction (FDR q < 0.05). ‘ns’ indicates statistical test did not survive FDR correction. We note that there was no statistical test between meta-matching finetune and classical transfer learning because the bootstrapping procedure was too expensive for the two methods. Colors indicate effect sizes of differences (Cohen’s D) between approaches. Light green color indicates effect size **≥** 0.8. Dark green color indicates 0 ≤ effect size < 0.8. Dark pink indicates ™0.8 < effect size < 0. Light pink color indicates effect size ≤ ™0.8. There is no color for the comparison between meta-matching finetune and stacking since they are both our proposed methods.

**Figure 8.**
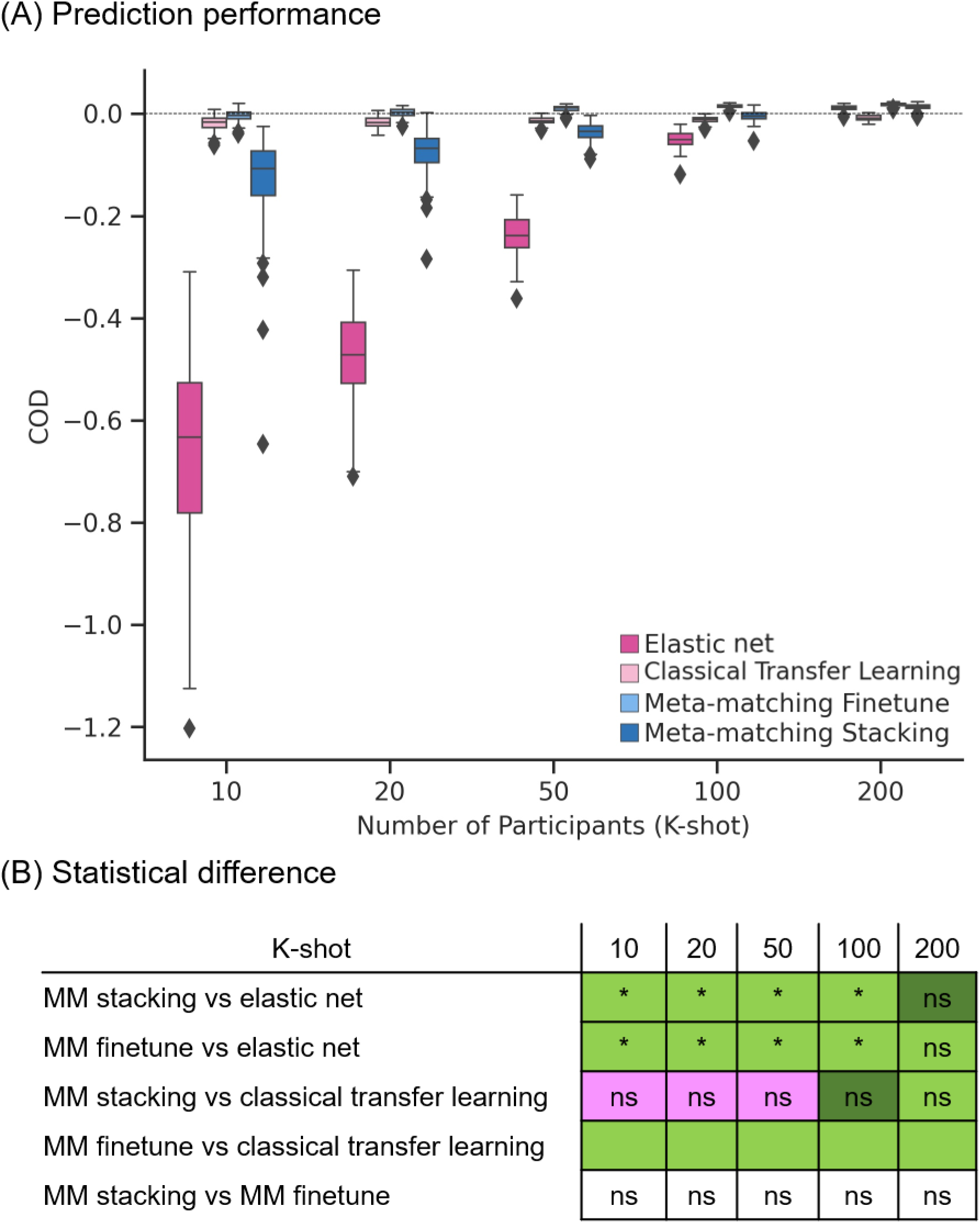
Meta-matching compared favorably with elastic net and direct transfer learning in terms of coefficient of determination (COD) when translating models from UK Biobank to HCP-YA dataset. (A) Phenotypic prediction performance (COD) averaged across 35 meta-test phenotypes in the HCP-YA dataset. X-axis is the number of participants from the HCP-YA dataset used to train an elastic net or adapt the pretrained model from the meta-training set. Each boxplot shows the distribution of performance over 100 repetitions of sampling K participants. (B) Statistical difference between the prediction performance (COD) of baseline methods and meta-matching algorithms. P values were calculated based on a two-sided bootstrapping test. ‘*’ indicates statistical significance after multiple comparisons correction (FDR q < 0.05). ‘ns’ indicates statistical test did not survive FDR correction. We note that there was no statistical test between meta-matching finetune and classical transfer learning because the bootstrapping procedure was too expensive for the two methods. Colors indicate effect sizes of differences (Cohen’s D) between approaches. Light green color indicates effect size **≥** 0.8. Dark green color indicates 0 ≤ effect size < 0.8. Dark pink indicates ™0.8 < effect size < 0. Light pink color indicates effect size ≤ ™0.8. There is no color for the comparison between meta-matching finetune and stacking since they are both our proposed methods.

In the case of Pearson’s correlation (Figure 7), both meta-matching finetune and meta-matching stacking were better than elastic net and classical transfer learning for all values of K with large effect sizes (light green in Figure 7B). Meta-matching finetune was statistically better than elastic net for K = 20 and 50. Meta-matching stacking was statistically better than classical transfer learning for K **≥** 20. For this cross-dataset analysis, meta-matching finetune was generally numerically better, but not statistically better than meta-matching stacking.

In the case of COD (Figure 8), both meta-matching finetune and meta-matching stacking greatly outperformed elastic net for K ≤ 100. For K ≤ 50, classical transfer learning was numerically better than meta-matching stacking with large effect sizes (light pink in Figure 8B), but the differences were not significant. From K ≥ 100, meta-matching stacking was numerically better, but not statistically better than transfer learning.

On the other hand, for all values of K, meta-matching finetune was numerically better than classical transfer learning with large effect sizes (light green in Figure 8B). We note that there was no statistical test between meta-matching finetune and classical transfer learning because of the huge computational cost of the two approaches, so no bootstrapping was performed for either approach.

Another relevant point is that COD for all approaches were negative (or almost zero) for K = 10, and only positive for meta-matching finetune for K ≥ 20, suggesting that absolute prediction accuracy (i.e., COD) is difficult even with meta-learning or transfer learning when the sample size is very small.

Overall, meta-matching compared favorably with respect to elastic net for all values of K for both evaluation metrics (Pearson’s correlation and COD). On the other hand, meta-matching compared favorably with respect to transfer learning for all values of K for Pearson’s correlation and for K ≥ 100 for COD.

### 3.3. Meta-matching outperforms baselines in the HCP-Aging dataset

Figures 9A and 10A show the Pearson’s correlation and COD respectively, averaged across all 45 meta-test phenotypes in the HCP-Aging dataset. Each boxplot represents 100 random samplings of K participants. Figures 9B and 10B show the outcomes of the statistical tests obtained by a bootstrapping procedure (Supplementary Methods S2). The actual p values are reported in Table S8. Colors indicate effect sizes of differences (Cohen’s D) between approaches.

**Figure 9.**
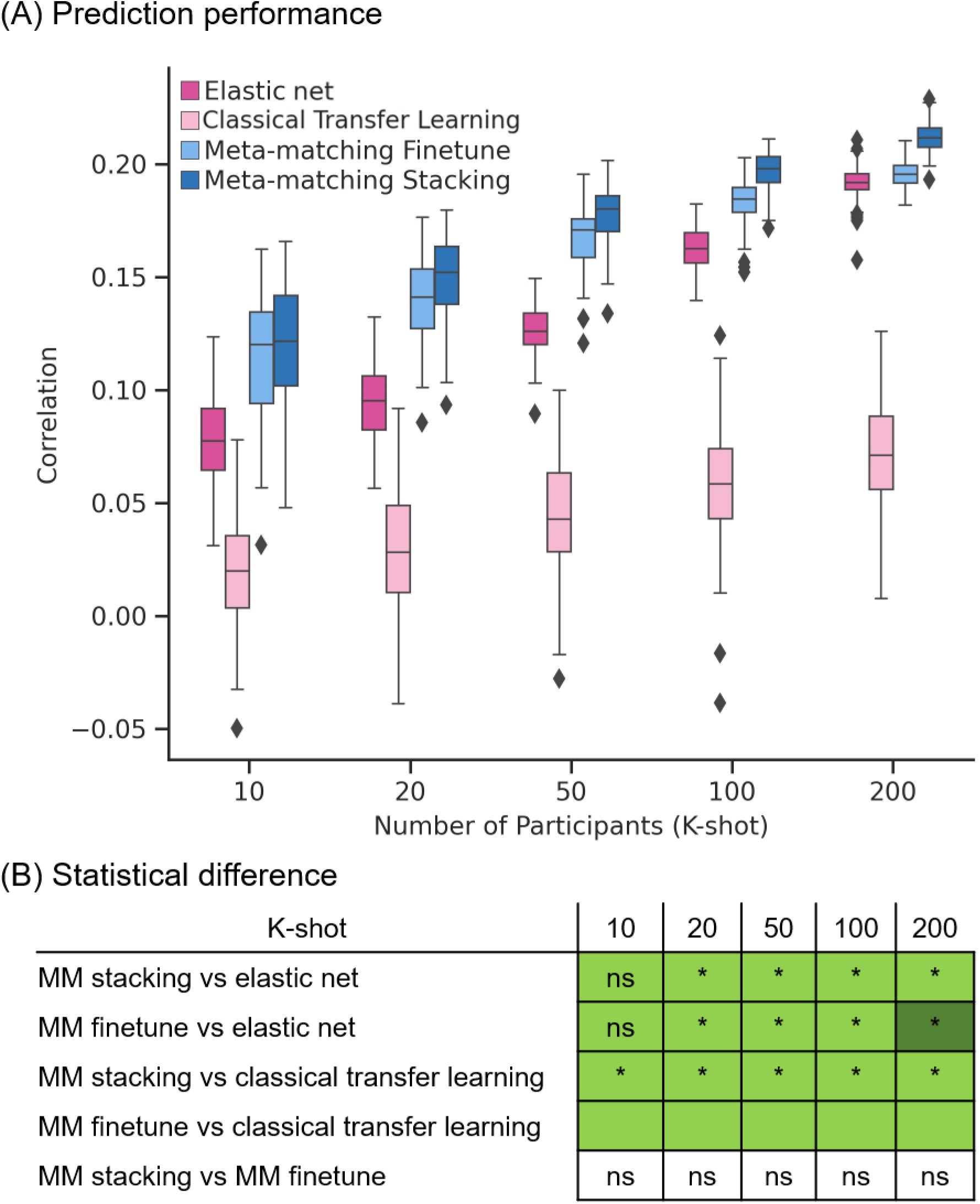
Meta-matching compared favorably with elastic net and direct transfer learning in terms of Pearson’s correlation when translating models from UK Biobank to HCP-Aging dataset. (A) Phenotypic prediction performance (Pearson’s correlation) averaged across 45 meta-test phenotypes in the HCP-Aging dataset. X-axis is the number of participants from the HCP-Aging dataset used to train an elastic net or adapt the pretrained model from the meta-training set. Each boxplot shows the distribution of performance over 100 repetitions of sampling K participants. (B) Statistical difference between the prediction performance (Pearson’s correlation) of baseline methods and meta-matching algorithms. P values were calculated based on a two-sided bootstrapping test. ‘*’ indicates statistical significance after multiple comparisons correction (FDR q < 0.05). ‘ns’ indicates statistical test did not survive FDR correction. We note that there was no statistical test between meta-matching finetune and classical transfer learning because the bootstrapping procedure was too expensive for the two methods. Colors indicate effect sizes of differences (Cohen’s D) between approaches. Light green color indicates effect size **≥** 0.8. Dark green color indicates 0 ≤ effect size < 0.8. Dark pink indicates ™0.8 < effect size < 0. Light pink color indicates effect size ≤ ™0.8. There is no color for the comparison between meta-matching finetune and stacking since they are both our proposed methods.

**Figure 10.**
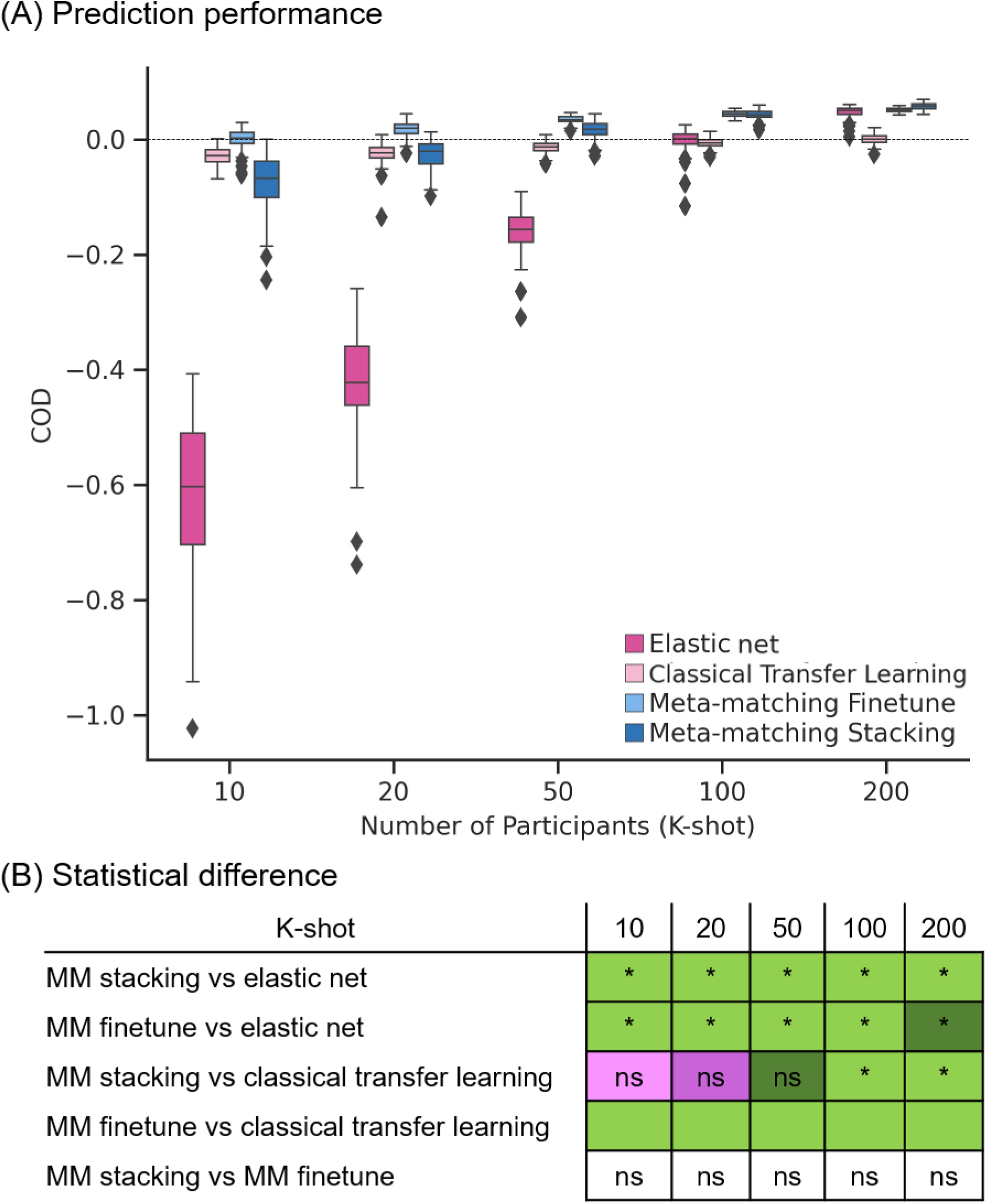
Meta-matching compared favorably with elastic net and direct transfer learning in terms of coefficient of determination (COD) when translating models from UK Biobank to HCP-Aging dataset. (A) Phenotypic prediction performance (COD) averaged across 45 meta-test phenotypes in the HCP-Aging dataset. X-axis is the number of participants from the HCP-Aging dataset used to train an elastic net or adapt the pretrained model from the meta-training set. Each boxplot shows the distribution of performance over 100 repetitions of sampling K participants. (B) Statistical difference between the prediction performance (COD) of baseline methods and meta-matching algorithms. P values were calculated based on a two-sided bootstrapping test. ‘*’ indicates statistical significance after multiple comparisons correction (FDR q < 0.05). ‘ns’ indicates statistical test did not survive FDR correction. We note that there was no statistical test between meta-matching finetune and classical transfer learning because the bootstrapping procedure was too expensive for the two methods. Colors indicate effect sizes of differences (Cohen’s D) between approaches. Light green color indicates effect size ≥ 0.8. Dark green color indicates 0 ≤ effect size < 0.8. Dark pink indicates ™0.8 < effect size < 0. Light pink color indicates effect size ≤ ™0.8. There is no color for the comparison between meta-matching finetune and stacking since they are both our proposed methods.

In the case of Pearson’s correlation (Figure 9), both meta-matching finetune and meta-matching stacking greatly outperformed elastic net and classical transfer learning for most values of K. Meta-matching stacking was statistically better than elastic net for K **≥** 20. Meta-matching stacking was statistically better than classical transfer learning for all values of K. For this cross-dataset analysis, meta-matching stacking was numerically better, but not statistically better than meta-matching finetune.

In the case of COD (Figure 10), both meta-matching finetune and meta-matching stacking greatly outperformed elastic net for all values of K. For K ≤ 20, classical transfer learning was numerically better, but not statistically better than meta-matching stacking. From K ≥ 50, meta-matching stacking was numerically better than transfer learning with statistical significance achieved for K ≥ 100.

On the other hand, for all values of K, meta-matching finetune was numerically better than classical transfer learning with large effect sizes (light green in Figure 10B). We note that there was no statistical test between meta-matching finetune and classical transfer learning because of the huge computational cost of the two approaches, so no bootstrapping was performed for either approach.

Another relevant point is that COD for all approaches were negative (or almost zero) for K = 10, and only positive for meta-matching finetune for K ≥ 20, suggesting that absolute prediction accuracy (i.e., COD) is difficult even with meta-learning or transfer learning when the sample size is very small.

Overall, meta-matching was better than elastic net for all values of K for both evaluation metrics (Pearson’s correlation and COD). On the other hand, meta-matching compared favorably with respect to transfer learning for all values of K for Pearson’s correlation and for K ≥ 50 for COD.

### 3.4. Different improvements on different phenotypes

Overall, meta-matching improved prediction on average across multiple phenotypes. However, we note that the improvement was not uniform across phenotypes. Figure 11 illustrates the prediction performance (Pearson’s correlation) of three non-brain-imaging phenotypes for K = 100 participants. In the case of the HCP-YA dataset (Figure 11A), meta-matching finetune compared favorably with other approaches for predicting dexterity and language, but only achieved similar prediction accuracy on emotion. In the case of HCP-Aging dataset (Figure 11B), meta-matching stacking compared favorably with other approaches for predicting fear somatic and anger aggression, but only achieved similar prediction accuracy on perceived rejection.

**Figure 11.**
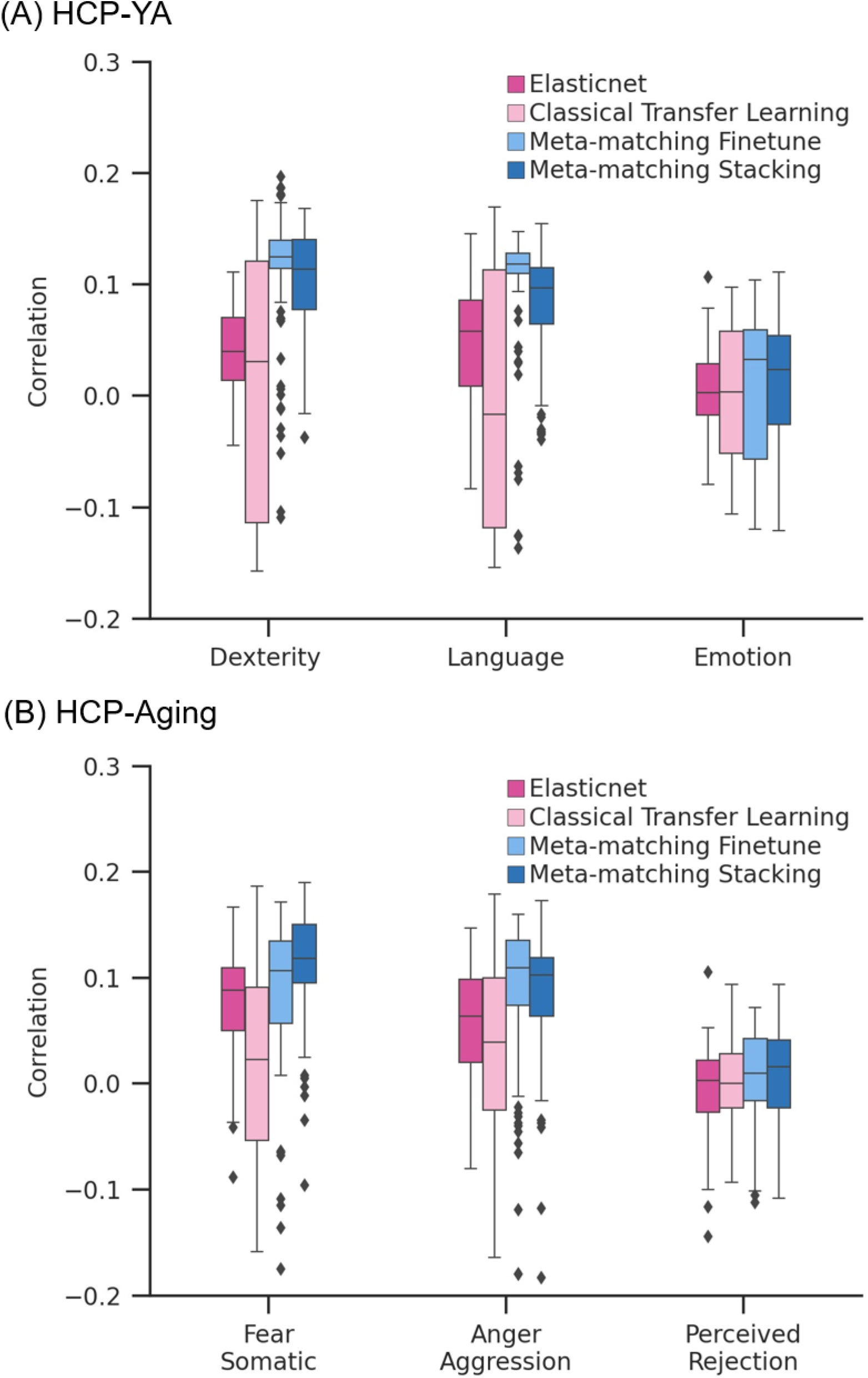
**Examples of prediction performance (Pearson’s correlation) for different non-brain-imaging phenotypes** in the (A) HCP-YA and (B) HCP-Aging datasets in the case of K = 100 participants.

Given that meta-matching exploits correlations among phenotypes, we hypothesized that variability in prediction improvements was driven by inter-phenotype correlations between the meta-training and meta-test sets. Figure 12 shows the performance improvement (Pearson’s correlation) of meta-matching stacking as a function of the maximum correlation between each meta-test phenotype and meta-training phenotype in the within-UK Biobank analysis. As expected, meta-test phenotypes with stronger correlations with at least one meta-training phenotype led to greater prediction improvement with meta-matching. We note that this analysis required meta-training and meta-test phenotypes to be present in the same participants, so could only be performed for the within-UK Biobank analysis.

**Figure 12.**
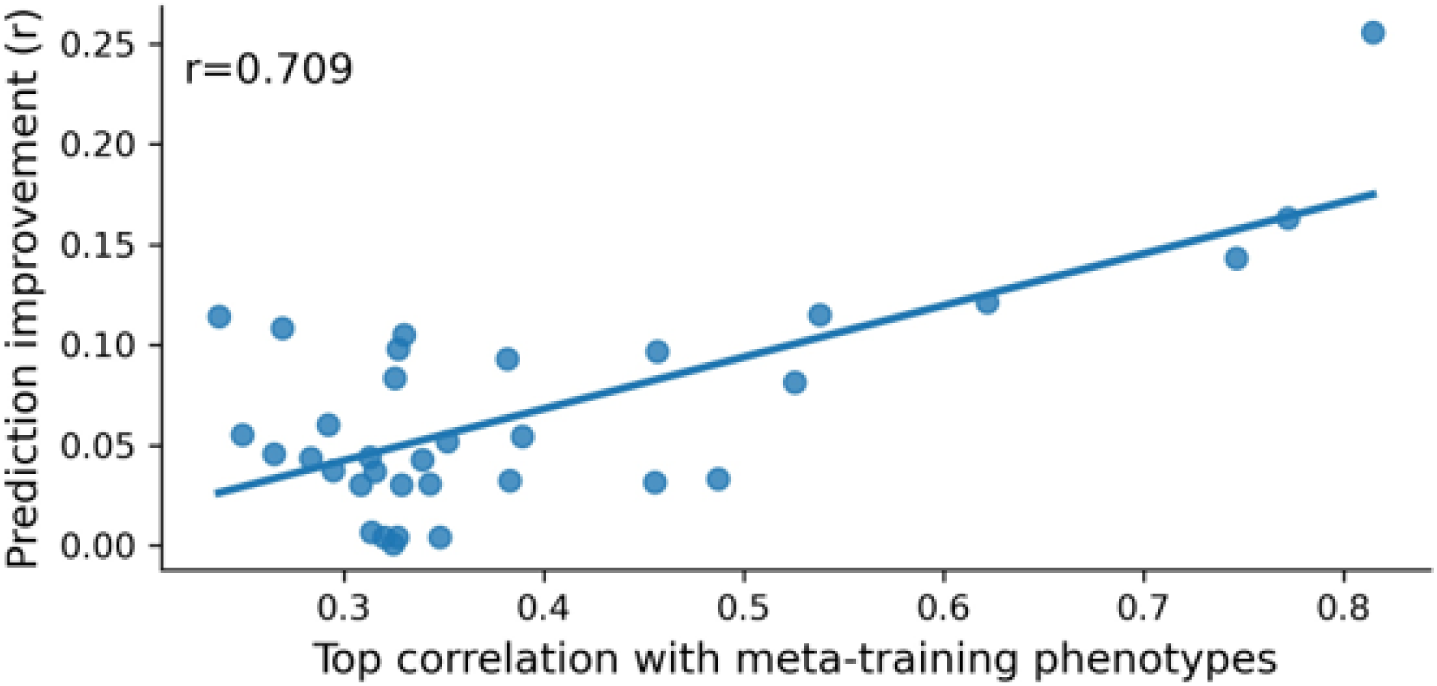
Prediction improvements were driven by correlations between meta-training and meta-test phenotypes. Vertical axis shows the prediction improvement of meta-matching stacking with respect to elastic net baseline under the 100-shot scenario. Prediction performance was measured using Pearson’s correlation. Each dot represents a meta-test phenotype. Horizontal axis shows each test phenotype’s top absolute Pearson’s correlation with phenotypes in the meta-training set. Test phenotypes with stronger correlations with at least one training phenotype led to greater prediction improvement with meta-matching.

### 3.5. Interpreting meta-matching stacking with Haufe transform

Figure 13 illustrates the feature importance maps obtained from the Haufe transform for predicting the Rey Auditory Verbal Learning Test (RAVLT) score and Montreal Cognitive Assessment (MOCA) in the HCP-Aging dataset for K = 100. We note that a higher RAVLT or MOCA scores indicated better cognition.

**Figure 13.**
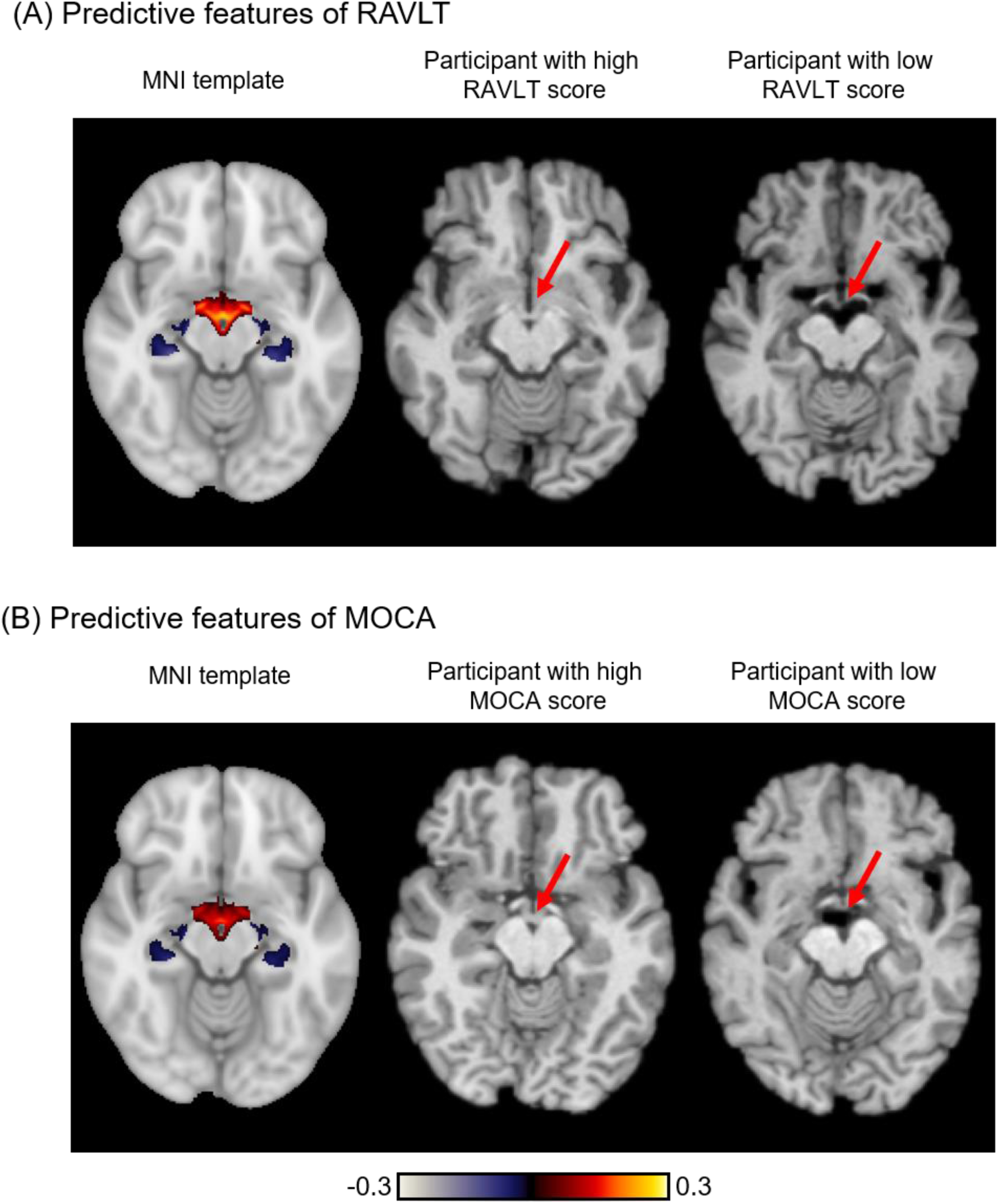
Feature importance of meta-matching stacking in the HCP-Aging dataset for K = 100 participants. (A) Feature importance map of meta-matching stacking from predicting Rey Auditory Verbal Learning Test (RAVLT) score. Left panel shows the feature importance map on the MNI152 template. A positive (or negative) feature importance value indicates that higher intensity was associated with predicting greater (or lower) phenotypic values. Middle panel shows an example participant with high RAVLT score. Right panel shows an example participant with low RAVLT score. (B) Feature importance map of meta-matching stacking from predicting the Montreal Cognitive Assessment (MOCA) score. Left panel shows the feature importance map on the MNI152 template. Middle panel shows an example participant with high MOCA score. Right panel shows an example participant with low MOCA score.

Since we are using T1 intensity for prediction, linking the feature importance values to the underlying biology needs to be done with care. For both RAVLT and MOCA, positive feature importance values were observed in the ventral diencephalon and the third ventricle (left panels of Figures 9A and 9B), which suggested that higher T1 value led to prediction of better cognition (higher RAVLT and MOCA scores). By observing participants who performed poorly (right panels of Figures 9A and 9B) and participants who performed well (middle panels of Figures 9A and 9B), we inferred that the prediction might be partially driven by enlarged ventricles in participants with worse cognition (arrows in Figure 9), yielding a lower T1 value in the region.

Similarly, we observed negative feature importance values at or near the left and right hippocampi, which suggested that higher T1 value led to prediction of worse cognition (lower RAVLT and MOCA scores). By observing participants who performed poorly (right panels of Figures 9A and 9B) and participants who performed well (middle panels of Figures 9A and 9B), we inferred that the prediction might be partially driven by gray matter loss at or near the hippocampi, yielding a higher T1 value in the region, consistent with the aging literature (Apostolova et al., 2012; Ritter et al., 2017).

## 4. Discussion

In this study, we adapted two meta-matching variants from our previous study (He et al., 2022) to translate prediction models trained from large-scale T1-weighted anatomical MRI datasets to predict new non-brain-imaging phenotypes in small-scale T1-weighted anatomical MRI datasets. We demonstrated that meta-matching finetune and meta-matching stacking greatly outperformed classical elastic net and classical transfer learning when the number of participants ≤ 200. Meta-matching performed well even when translating from a large-scale dataset (UK Biobank) to a small dataset (HCP-YA or HCP-Aging) with different scanners, acquisition, demographics, and pre-processing.

Across all analyses in the UK Biobank, HCP-YA and HCP-Aging datasets (Figures 5 to 10), meta-matching consistently outperformed elastic net across both evaluation metrics (correlation and COD). It is worth noting that the elastic net utilized thickness and volumetric measures generated by FreeSurfer, instead of the intensity values of T1 images (like meta-matching and transfer learning). Given that we are working in the small sample regime with K ≤ 200 training participants, we believe that the small number of less than 200 pre-defined morphometric features together with elastic net provides a powerful classical baseline.

When using Pearson’s correlation as an evaluation metric, transfer learning performed poorly with substantially worse performance than both meta-matching variants and even elastic net (Figures 5, 7 and 9). On the other hand, when using COD as an evaluation metric, transfer learning was more competitive with respect to the other approaches (Figures 6, 8 and 10). More specifically, transfer learning was numerically better (but not statistically better) than meta-matching stacking for small values of K, while meta-matching stacking was numerically better (and sometimes statistically better) than transfer learning for larger values of K.

On the other hand, meta-matching finetune outperformed transfer learning for most values of K even in the case of COD. We note that meta-matching finetune is similar to classical transfer learning in the sense that the last two layers of the CNN were finetuned. However, while transfer learning initialized the last layer of the CNN from scratch (Section 2.3.2), meta-matching finetune retained the weights leading to the output node that predicted the K meta-test participants the best (for each meta-test phenotype). This further supported the importance of the meta-matching approach.

Overall, meta-matching stacking was the best for the Pearson’s correlation metric, while meta-matching finetune was the best for COD. Pearson’s correlation is a measure of relative prediction performance, while COD is a measure of absolute prediction performance (Finn et al., 2015; Scheinost et al., 2019; Poldrack et al., 2020). Therefore, researchers more focused on relative prediction performance might consider using meta-matching stacking, while researchers more focused on absolute prediction performance might consider using meta-matching finetune. Furthermore, all approaches achieved negative or close to zero COD when K ≤ 20, suggesting that absolute prediction remains out of reach in the very small sample regime.

Meta-matching models can be interpreted at the level of imaging features by using the Haufe transform (Haufe et al., 2014). To illustrate this procedure, we applied the Haufe transform (Haufe et al., 2014) to the translated meta-matching stacking models in the HCP-Aging dataset (Figure 13). For a given meta-test phenotype, Haufe transform was calculated as the covariance between the phenotype’s prediction based on the meta-matching stacking model and the intensity value of each T1 voxel (across K participants), yielding a 3D volume. We found that poorer cognitive performance in terms of worse RAVLT and MOCA scores were related to greater gray matter atrophy and larger ventricular size, which is consistent with the aging literature (Apostolova et al., 2012; Ritter et al., 2017). Meta-matching finetune can be interpreted in a similar fashion.

In addition to interpreting meta-matching models at the level of brain-imaging features, the meta-matching models can also be interpreted at the level of phenotypic traits. In the case of meta-matching stacking, this can again be achieved using the Haufe transform. To illustrate this, let us consider the pretrained 3D CNN model from the UK Biobank with 67 prediction outputs. This 3D CNN model can be translated to predict a new meta-test phenotype using K participants from the meta-test set using the stacking procedure. The Haufe transform can then be calculated as the covariance (across the K participants) between the phenotype’s prediction from the final stacking model and the 67 inputs to the stacking model, yielding a vector of length 67, which indicates the relative importance of the original 67 meta-training phenotypes for predicting the meta-test phenotype.

Because meta-matching exploits correlations between the phenotypes of meta-training and meta-test sets, the amount of prediction improvement strongly relied on the strongest correlations between the meta-test phenotype and meta-training phenotypes (Figure 12). Consequently, not all phenotypes might benefit from meta-matching. However, we note that this limitation exists for all meta-learning and transfer learning algorithms – model transfer is easier if the source and target domains are more similar; performance will degrade if source and target domains are very different.

In future work, we seek to aggregate more and diverse large-scale population-level datasets targeting different populations, including children, lifespan and a variety of disorders. These additional datasets will not only provide a larger diversity of phenotypic measures but will also allow greater sampling of different MRI scanners, acquisition parameters and demographics. By pretraining on a wider range of datasets, we hope to see further improvements in the meta-matching approaches.

## 5. Conclusion

In this study, we showed that meta-matching can be used to translate T1-based phenotypic prediction models from large source datasets to predict new phenotypes in small target datasets. By exploiting correlations between phenotypes, meta-matching greatly outperformed elastic net and classical transfer learning, both when translating models within the same dataset, as well as translating models across datasets with different MRI scanners, acquisition protocols and demographics. Overall, our results demonstrated the versatility of the meta-matching framework.

## 6. Acknowledgements

Our research is supported by the NUS Yong Loo Lin School of Medicine (NUHSRO/2020/124/TMR/LOA), the Singapore National Medical Research Council (NMRC) LCG (OFLCG19May-0035), NMRC CTG-IIT (CTGIIT23jan-0001), NMRC STaR (STaR20nov-0003), Singapore Ministry of Health (MOH) Centre Grant (CG21APR1009), the Temasek Foundation (TF2223-IMH-01), and the United States National Institutes of Health (R01MH120080 & R01MH133334). Our computational work was partially performed on resources of the National Supercomputing Centre, Singapore (https://www.nscc.sg). Any opinions, findings and conclusions or recommendations expressed in this material are those of the authors and do not reflect the views of the Singapore NRF, NMRC or MOH. This research has been conducted using the UK Biobank Resource under application number 25163, and the Human Connectome Project, WU-Minn Consortium (Principal Investigators: David Van Essen and Kamil Ugurbil; 1U54MH091657) funded by the 16 NIH Institutes and Centers that support the NIH Blueprint for Neuroscience Research, and the National Institute On Aging of the National Institutes of Health under Award Number U01AG052564 and by funds provided by the McDonnell Center for Systems Neuroscience at Washington University in St. Louis. The HCP-Aging 2.0 Release data used in this report came from DOI: 10.15154/1520707.; and by the McDonnell Center for Systems Neuroscience at Washington University.

## Author Contribution

N.W., L.A., C.Z., R.K., P.C., D.B., S.B.E., A.J.H. and B.T.T.Y. designed the research. N.W., conducted the research. N.W., L.A., C.Z., R.K., P.C., D.B., S.B.E., A.J.H. and B.T.T.Y. interpreted the results. N.W.and B.T.T.Y. wrote the manuscript and made the figures. N.W., L.A., and C.Z. reviewed and published the code. All authors contributed to project direction via discussion. All authors edited the manuscript.

