## Supplementary Materials for "Translating phenotypic prediction models from big to small anatomical MRI data using meta-matching"

This supplementary material is divided into Supplementary Methods and Supplementary Results to complement the Methods and Results sections in the main text, respectively.

#### Supplemental Methods

##### S1. Details of pre-training 3D-CNN on the UK Biobank

Details about the 3D CNN model (pre-trained on the UK Biobank) are shown below. We note that hyperparameters were empirically determined based on the meta-training set from the within-UK Biobank analysis (Figure 1A). Both within-UK Biobank and cross-dataset analyses used the same set of hyperparameters (Table S1).

During training (as well as when performing meta-matching finetune and transfer learning), mean square error was used as the loss function. The output layer of the 3D CNN model had 33 nodes for within UK Biobank analyses and 67 nodes for cross-dataset analyses. The maximum number of training epochs was 100 for training in the meta-training set. The initial learning rate was 1e-5 and decreased by a factor 10 after every 50 epochs. The meta-training validation set was used to evaluate the validation loss for early stopping.

The 3D CNN model was implemented using PyTorch (Paszke et al., 2017).

| Hyperparameter | Value |
|---|---|
| Number of 3D convolutional blocks | 6 |
| Number of channels | [32 64 128 256 256 64] |
| Dropout rate | 0.1 |
| Starting learning rate | 1e-5 |
| Weight decay rate | 0.02 |
| Batch size | 8 |

**Table S1.** Hyperparameters for the 3D CNN model.

#### S2. Bootstrapping procedure

Following our previous study (He et al., 2022), we used a bootstrapping procedure to evaluate whether there is a statistical difference between algorithms. The bootstrap sampling was repeated 1,000 times. For each bootstrap sampling, we randomly chose K participants in the meta-test set with replacement, and the remaining participants in the meta-test set were used as test participants. We applied elastic net and meta-matching stacking separately on each of the 1,000 bootstrapped samples, resulting in 1,000 COD and Pearson's correlation from the bootstrapped samples for each method. 1,000 bootstrapping was not performed for classical transfer learning and meta-matching finetune because it would have required 40 days of computing time (on a single GPU).

We calculated the statistical significance for Pearson's correlation and COD separately. For ease of illustration, we focused on COD to explain the procedure for the statistical test. To compute the statistical difference between two approaches with 1,000 bootstrapped samples (i.e., elastic net and meta-matching stacking), we fitted a Gaussian distribution to the difference of 1,000 COD between the two algorithms. We then calculated the p-value as follows:

$$p = \begin{cases} 2 \times \mathrm{CDF}(0) & \text{if } \mu \geq 0 \\ 2 \times \left(1 - \mathrm{CDF}(0)\right) & \text{if } \mu < 0 \end{cases} \tag{1}$$

CDF and $\mu$ are the cumulative distribution function and the mean of the Gaussian distribution separately.

To compute the statistical difference between one approaches with 1,000 bootstrapped samples and another approach with no bootstrapped samples (e.g., elastic net versus meta-matching finetune), we fitted a Gaussian distribution to the 1,000 COD of elastic net. We then calculated the p-value as follows:

$$p = \begin{cases} 2 \times \mathrm{CDF}(u_m) & \text{if } \mu \geq u_m \\ 2 \times \left(1 - \mathrm{CDF}(u_m)\right) & \text{if } \mu < u_m \end{cases} \tag{2}$$

CDF and $\mu$ are the cumulative distribution function and the mean of the Gaussian distribution separately. $u_m$ is the average COD of meta-matching finetune across the 100 random repeats of K-shot. The procedure for Pearson's correlation is the same as COD. FDR ($q < 0.05$) is used for multiple comparisons correction.

| UKBB field | Description |
|---|---|
| #household | number of people in household |
| #Mem C1 | numeric memory principal component 1 |
| Age | age |
| Age edu | age completed full time education |
| Alcohol 1 | average monthly spirits intake |
| Alcohol 2 | average weekly champagne plus white wine intake |
| Alcohol 3 | average weekly beer plus cider intake |
| Blood C2 | blood assays principal component 2 |
| Blood C3 | blood assays principal component 3 |
| Blood C4 | blood assays principal component 4 |
| Blood C5 | blood assays principal component 5 |
| Body C1 | anthropometry principal component 1 |
| Body C2 | anthropometry principal component 2 |
| Body C3 | anthropometry principal component 3 |
| Bone C1 | bone-densitometry of heel principal component 1 |
| Bone C3 | bone-densitometry of heel principal component 3 |
| BP eye C2 | blood pressure & eye measures component 2 |
| BP eye C3 | blood pressure & eye measures principal component 3 |
| BP eye C4 | blood pressure & eye measures principal component 4 |
| BP eye C5 | blood pressure & eye measures principal component 5 |
| BP eye C6 | blood pressure & eye measures principal component 6 |
| Breath C1 | spirometry principal component 1 |
| Cancer C1 | cancer principal component 1 |
| Carotid C1 | carotid ultrasound principal component 1 |
| Carotid C5 | carotid ultrasound principal component 5 |
| Deprive C1 | multiple deprivation principal component 1 |
| Digit 1 | symbol digit substitution principal component 1 |
| Digit-o C1 | symbol digit substitution online principal component 1 |
| Digit-o C6 | symbol digit substitution online principal component 6 |
| Dur C1 | process durations principal component 1 |
| Dur C2 | process durations principal component 2 |
| Dur C4 | process durations principal component 4 |

| | |
|---|---|
| ECG C1 | ECG measures principal component 1 |
| ECG C2 | ECG measures principal component 2 |
| ECG C3 | ecg measures principal component 3 |
| ECG C6 | ECG measures principal component 6 |
| Family C1 | family history (parent's age) principal component 1 |
| Fluid Int. | fluid intelligence |
| Genetic C1 | genetic principal components and heterozygosity principal component 1 |
| Grip C1 | hand grip strength principal component 1 |
| Hearing | hearing signal-to-noise-ratio (snr) of triplet (left) |
| Illness C1 | non-cancer illness principal component 1 |
| Illness C4 | non-cancer illness principal component 4 |
| Loc C1 | location principal component 1 |
| Match | pairs matching |
| Match-o | pairs matching online |
| Matrix C1 | matrix pattern completion principal component 1 |
| Matrix C2 | matrix pattern completion principal component 2 |
| Matrix C3 | matrix pattern completion principal component 3 |
| Neuro | neuroticism score |
| ProMem C1 | prospective memory principal component 1 |
| RT C1 | reaction time principal component 1 |
| Sex | sex |
| Sex G C1 | genotype sex inference principal component 1 |
| Sex G C2 | genotype sex inference principal component 2 |
| Sleep | sleep duration per day |
| Smoke C1 | smoke principal component 1 |
| Time drive | time spent driving per day |
| Time TV | time spent watching television (tv) per day |
| Time walk | number of days walked 10+ minutes per week |
| Tower C1 | tower rearranging principal component 1 |
| Trail C1 | trail making principal component 1 |
| Trail-o C1 | trail making online principal component 1 |
| Trail-o C4 | trail making online principal component 4 |
| Travel | frequency of travelling from home to job workplace per week |
| Urine C1 | urine assays principal component 1 |

| | |
|---|---|
| Work | weekly length of working hour for main job |

**Table S1.** Dictionary of 67 phenotypes in UK Biobank.

| HCP field | Description |
|---|---|
| PicSeq_Unadj | Visual Episodic Memory |
| CardSort_Unadj | Cognitive Flexibility (DCCS) |
| Flanker_Unadj | Inhibition (Flanker Task) |
| PMAT24_A_CR | Fluid Intelligence (PMAT) |
| ReadEng_Unadj | Vocabulary (Pronunciation) |
| PicVocab_Unadj | Vocabulary (Picture Matching) |
| ProcSpeed_Unadj | Processing Speed |
| DDic_AUC_40K | Delay Discounting |
| VSPLOT_TC | Spatial Orientation |
| SCPT_SPEC | Sustained Attention – Spec. |
| ListSort_Unadj | Working Memory (List Sorting) |
| MMSE_Score | Cognitive Status (MMSE) |
| PSQI_Score | Sleep Quality (PSQI) |
| Endurance_Unadj | Walking Endurance |
| GaitSpeed_Unadj | Walking Speed |
| Dexterity_Unadj | Manual Dexterity |
| Strength_Unadj | Grip Strength |
| Taste_Unadj | Taste Intensity |
| Emotion_Task_Face_Acc | Emotional Face Matching |
| Language_Task_Math_Avg_Difficulty_Level | Arithmetic |
| Language_Task_Story_Avg_Difficulty_Level | Story Comprehension |
| Relational_Task_Acc | Relational Processing |
| WM_Task_Acc | Working Memory (N-back) |
| NEOFAC_A | Agreeableness (NEO) |
| NEOFAC_O | Openness (NEO) |
| NEOFAC_C | Conscientiousness (NEO) |
| NEOFAC_E | Extraversion (NEO) |
| AngAggr_Unadj | Anger – Aggression |
| FearAffect_Unadj | Fear – Affect |
| Sadness_Unadj | Sadness |
| LifeSatisf_Unadj | Life Satisfaction |

| | |
|---|---|
| MeanPurp_Unadj | Meaning & Purpose |
| Loneliness_Unadj | Loneliness |
| PercStress_Unadj | Perceived Stress |
| SelfEff_Unadj | Self-Efficacy |

**Table S2.** Dictionary of 35 phenotypes in the HCP-YA dataset.

| HCP-Aging field | Description |
|---|---|
| LifeSatisf | Life Satisfaction |
| MeanPurp | Meaning & Purpose |
| PosAffect | Positive Affect |
| Sadness | Sadness |
| FearAffect | Fear – Affect |
| FearSomat | Fear – Somatic |
| AngAffect | Anger – Affect |
| AngHostil | Anger – Hostility |
| AngAggr | Anger – Aggression |
| ER40ANG | ER40 Correct Anger Identifications |
| ER40FEAR | ER40 Correct Fear Identifications |
| ER40NOE | ER40 Correct No Emotion Identifications |
| ER40SAD | ER40 Correct Sad Identifications |
| EmotSupp | Emotion Support |
| InstruSupp | Instrumental Support |
| Friendship | Friendship |
| Loneliness | Loneliness |
| PercReject | Perceived Rejection |
| PercHostil | Perceived Hostility |
| PercStress | Perceived Stress |
| SelfEff | Self-Efficacy |
| NEOFAC_A | Agreeableness (NEO) |
| NEOFAC_O | Openness (NEO) |
| NEOFAC_C | Conscientiousness (NEO) |
| NEOFAC_N | Neuroticism (NEO) |
| NEOFAC_E | Extraversion (NEO) |
| Endurance | Walking Endurance |

| | |
|---|---|
| GaitSpeed | Walking Speed |
| Strength | Grip Strength |
| PicSeq | Visual Episodic Memory |
| CardSort | Cognitive Flexibility (DCCS) |
| Flanker | Inhibition (Flanker Task) |
| ReadEng | Vocabulary (Pronunciation) |
| PicVocab | Vocabulary (Picture Matching) |
| ProcSpeed | Processing Speed |
| DDisc_AUC_40K | Delay Discounting |
| ListSort | Working Memory (List Sorting) |
| raw_vat | Raw Score for the Visual Acuity |
| raw_pain | Raw Score for the Pain Interference |
| winright_ncorr | Words in Noise Number Correct Right |
| psqi_total | Sleep Quality (PSQI) |
| tmta_raw | Trailmaking test part A: Time to completion |
| tmtb_raw | Trailmaking test Part B: Time to completion |
| pea_ravlt_sd_tc | RAVLT Short Delay Total Correct |
| moca_total | Montreal Cognitive Assessment Total Score |

**Table S3.** Dictionary of 45 phenotypes on HCP-Aging dataset.

| brain ROI thickness | brain ROI thickness |
|---|---|
| lh_caudalanteriorcingulate_thickness | rh_caudalanteriorcingulate_thickness |
| lh_caudalmiddlefrontal_thickness | rh_caudalmiddlefrontal_thickness |
| lh_cuneus_thickness | rh_cuneus_thickness |
| lh_entorhinal_thickness | rh_entorhinal_thickness |
| lh_fusiform_thickness | rh_fusiform_thickness |
| lh_inferiorparietal_thickness | rh_inferiorparietal_thickness |
| lh_inferiortemporal_thickness | rh_inferiortemporal_thickness |
| lh_insula_thickness | rh_insula_thickness |
| lh_isthmuscingulate_thickness | rh_isthmuscingulate_thickness |
| lh_lateraloccipital_thickness | rh_lateraloccipital_thickness |
| lh_lateralorbitofrontal_thickness | rh_lateralorbitofrontal_thickness |
| lh_lingual_thickness | rh_lingual_thickness |
| lh_medialorbitofrontal_thickness | rh_medialorbitofrontal_thickness |

| | |
|---|---|
| lh_middletemporal_thickness | rh_middletemporal_thickness |
| lh_paracentral_thickness | rh_paracentral_thickness |
| lh_parahippocampal_thickness | rh_parahippocampal_thickness |
| lh_parsopercularis_thickness | rh_parsopercularis_thickness |
| lh_parsorbitalis_thickness | rh_parsorbitalis_thickness |
| lh_parstriangularis_thickness | rh_parstriangularis_thickness |
| lh_pericalcarine_thickness | rh_pericalcarine_thickness |
| lh_postcentral_thickness | rh_postcentral_thickness |
| lh_posteriorcingulate_thickness | rh_posteriorcingulate_thickness |
| lh_precentral_thickness | rh_precentral_thickness |
| lh_precuneus_thickness | rh_precuneus_thickness |
| lh_rostralanteriorcingulate_thickness | rh_rostralanteriorcingulate_thickness |
| lh_rostralmiddlefrontal_thickness | rh_rostralmiddlefrontal_thickness |
| lh_superiorfrontal_thickness | rh_superiorfrontal_thickness |
| lh_superiorparietal_thickness | rh_superiorparietal_thickness |
| lh_superiortemporal_thickness | rh_superiortemporal_thickness |
| lh_supramarginal_thickness | rh_supramarginal_thickness |
| lh_transversetemporal_thickness | rh_transversetemporal_thickness |

**Table S4.** Selected 62 brain ROI thicknesses for DKT40 atlas.

| **brain ROI volume** | **brain ROI volume** |
|---|---|
| lh_caudalanteriorcingulate_volume | rh_postcentral_volume |
| lh_caudalmiddlefrontal_volume | rh_posteriorcingulate_volume |
| lh_cuneus_volume | rh_precentral_volume |
| lh_entorhinal_volume | rh_precuneus_volume |
| lh_fusiform_volume | rh_rostralanteriorcingulate_volume |
| lh_inferiorparietal_volume | rh_rostralmiddlefrontal_volume |
| lh_inferiortemporal_volume | rh_superiorfrontal_volume |
| lh_insula_volume | rh_superiorparietal_volume |
| lh_isthmuscingulate_volume | rh_superiortemporal_volume |
| lh_lateraloccipital_volume | rh_supramarginal_volume |
| lh_lateralorbitofrontal_volume | rh_transversetemporal_volume |
| lh_lingual_volume | 3rd-Ventricle |
| lh_medialorbitofrontal_volume | 4th-Ventricle |
| lh_middletemporal_volume | Left-Accumbens-area |

| | |
|---|---|
| lh_paracentral_volume | Right-Accumbens-area |
| lh_parahippocampal_volume | Left-Amygdala |
| lh_parsopercularis_volume | Right-Amygdala |
| lh_parsorbitalis_volume | Brain-Stem |
| lh_parstriangularis_volume | CC_Anterior |
| lh_pericalcarine_volume | CC_Central |
| lh_postcentral_volume | CC_Mid_Anterior |
| lh_posteriorcingulate_volume | CC_Mid_Posterior |
| lh_precentral_volume | CC_Posterior |
| lh_precuneus_volume | CSF |
| lh_rostralanteriorcingulate_volume | Left-Caudate |
| lh_rostralmiddlefrontal_volume | Right-Caudate |
| lh_superiorfrontal_volume | Left-Cerebellum-Cortex |
| lh_superiorparietal_volume | Right-Cerebellum-Cortex |
| lh_superiortemporal_volume | Left-Cerebellum-White-Matter |
| lh_supramarginal_volume | Right-Cerebellum-White-Matter |
| lh_transversetemporal_volume | EstimatedTotalIntraCranialVol |
| rh_caudalanteriorcingulate_volume | Left-Hippocampus |
| rh_caudalmiddlefrontal_volume | Right-Hippocampus |
| rh_cuneus_volume | Left-Inf-Lat-Vent |
| rh_entorhinal_volume | Right-Inf-Lat-Vent |
| rh_fusiform_volume | Left-Lateral-Ventricle |
| rh_inferiorparietal_volume | Right-Lateral-Ventricle |
| rh_inferiortemporal_volume | Optic-Chiasm |
| rh_insula_volume | Left-Pallidum |
| rh_isthmuscingulate_volume | Right-Pallidum |
| rh_lateraloccipital_volume | Left-Putamen |
| rh_lateralorbitofrontal_volume | Right-Putamen |
| rh_lingual_volume | Left-Thalamus-Proper |
| rh_medialorbitofrontal_volume | Right-Thalamus-Proper |
| rh_middletemporal_volume | Left-VentralDC |
| rh_paracentral_volume | Right-VentralDC |
| rh_parahippocampal_volume | WM-hypointensities |
| rh_parsopercularis_volume | Left-choroid-plexus |
| rh_parsorbitalis_volume | Right-choroid-plexus |
| rh_parstriangularis_volume | Left-vessel |

| | |
|---|---|
| rh_pericalcarine_volume | Right-vessel |

**Table S5.** Selected 104 brain ROI volumes from DKT40 and ASEG atlases.

### Supplemental Results

| p-value | | K=10 | K=20 | K=50 | K=100 | K=200 |
|---|---|---|---|---|---|---|
| Correlation | MM stacking v.s. Elastic net | **7.17E-05** | **9.60E-08** | **2.78E-15** | **0** | **0** |
| | MM finetune v.s. Elastic net | **0.000109** | **3.07E-07** | **6.66E-16** | **0** | **1.55E-15** |
| | MM stacking v.s. Classical transfer learning | **1.62E-08** | **0** | **0** | **0** | **0** |
| | MM stacking v.s. MM finetune | 0.3695 | 0.2112 | 0.1415 | **0.0285** | **0.0003** |
| COD | MM stacking v.s. Elastic net | **0.0005** | **1.89E-06** | **1.78E-08** | **1.49E-06** | **6.65E-05** |
| | MM finetune v.s. Elastic net | **0.0001** | **4.68E-06** | **7.97E-08** | **9.02E-06** | **0.0031** |
| | MM stacking v.s. Classical transfer learning | 0.0962 | 0.5311 | 0.1315 | **4.85E-10** | **0** |
| | MM stacking v.s. MM finetune | 0.1337 | 0.1680 | 0.1539 | 0.6837 | 0.0533 |

**Table S6**. Meta-matching outperformed elastic net and classical transfer learning in UK Biobank. Statistical differences among the different algorithms for both correlation and COD metrics. P values were calculated based on a bootstrapping procedure. 'MM' stands for 'meta-matching'. Bold denotes statistical difference after correction for multiple comparisons (FDR $q < 0.05$).

| p-value | | K=10 | K=20 | K=50 | K=100 | K=200 |
|---|---|---|---|---|---|---|
| **Correlation** | MM stacking v.s. Elastic net | 0.1561 | 0.0990 | 0.0650 | 0.1499 | 0.3829 |
| | MM finetune v.s. Elastic net | 0.0358 | **0.0102** | **0.0051** | 0.0337 | 0.2024 |
| | MM stacking v.s. Classical transfer learning | 0.0392 | **0.0022** | **1.11E-07** | **0** | **0** |
| | MM stacking v.s. MM finetune | 0.9631 | 0.7811 | 0.4498 | 0.2682 | 0.3337 |
| **COD** | MM stacking v.s. Elastic net | **0.0151** | **0.0011** | **0.0010** | **0.0265** | 0.2798 |
| | MM finetune v.s. Elastic net | **0.0027** | **0.0001** | **5.70E-05** | **0.0012** | 0.0557 |
| | MM stacking v.s. Classical transfer learning | 0.1799 | 0.1146 | 0.1874 | 0.9794 | 0.1431 |
| | MM stacking v.s. MM finetune | 0.1324 | 0.0384 | **0.01226** | 0.0300 | 0.1109 |

**Table S7**. Meta-matching outperformed elastic net and classical transfer learning in HCP-YA. Statistical differences among the different algorithms for both correlation and COD metrics. P values were calculated based on a bootstrapping procedure. ‘MM’ stands for ‘meta-matching’. Bold denotes statistical difference after correction for multiple comparisons (FDR q < 0.05).

| p-value | | K=10 | K=20 | K=50 | K=100 | K=200 |
|---|---|---|---|---|---|---|
| **Correlation** | MM stacking v.s. Elastic net | 0.0304 | **0.0013** | **0.0001** | **0.0004** | **0.0043** |
| | MM finetune v.s. Elastic net | 0.0451 | **0.0035** | **0.0006** | **0.0025** | **0.0115** |
| | MM stacking v.s. Classical transfer learning | **9.16E-05** | **3.13E-09** | **0** | **0** | **0** |
| | MM stacking v.s. MM finetune | 0.8949 | 0.7928 | 0.6398 | 0.4795 | 0.4527 |
| **COD** | MM stacking v.s. Elastic net | **0.0047** | **2.84E-07** | **1.39E-06** | **0.0036** | **0.0116** |
| | MM finetune v.s. Elastic net | **0.0019** | **8.66E-08** | **7.20E-07** | **0.0023** | **0.0112** |
| | MM stacking v.s. Classical transfer learning | 0.3717 | 0.7631 | 0.2004 | **0.0009** | **1.25E-07** |
| | MM stacking v.s. MM finetune | 0.1712 | 0.0997 | 0.1387 | 0.3173 | 0.5261 |

**Table S8**. Meta-matching outperformed elastic net and classical transfer learning in HCP-Aging. Statistical differences among the different algorithms for both correlation and COD metrics. P values were calculated based on a bootstrapping procedure. 'MM' stands for 'meta-matching'. Bold denotes statistical difference after correction for multiple comparisons (FDR q < 0.05).

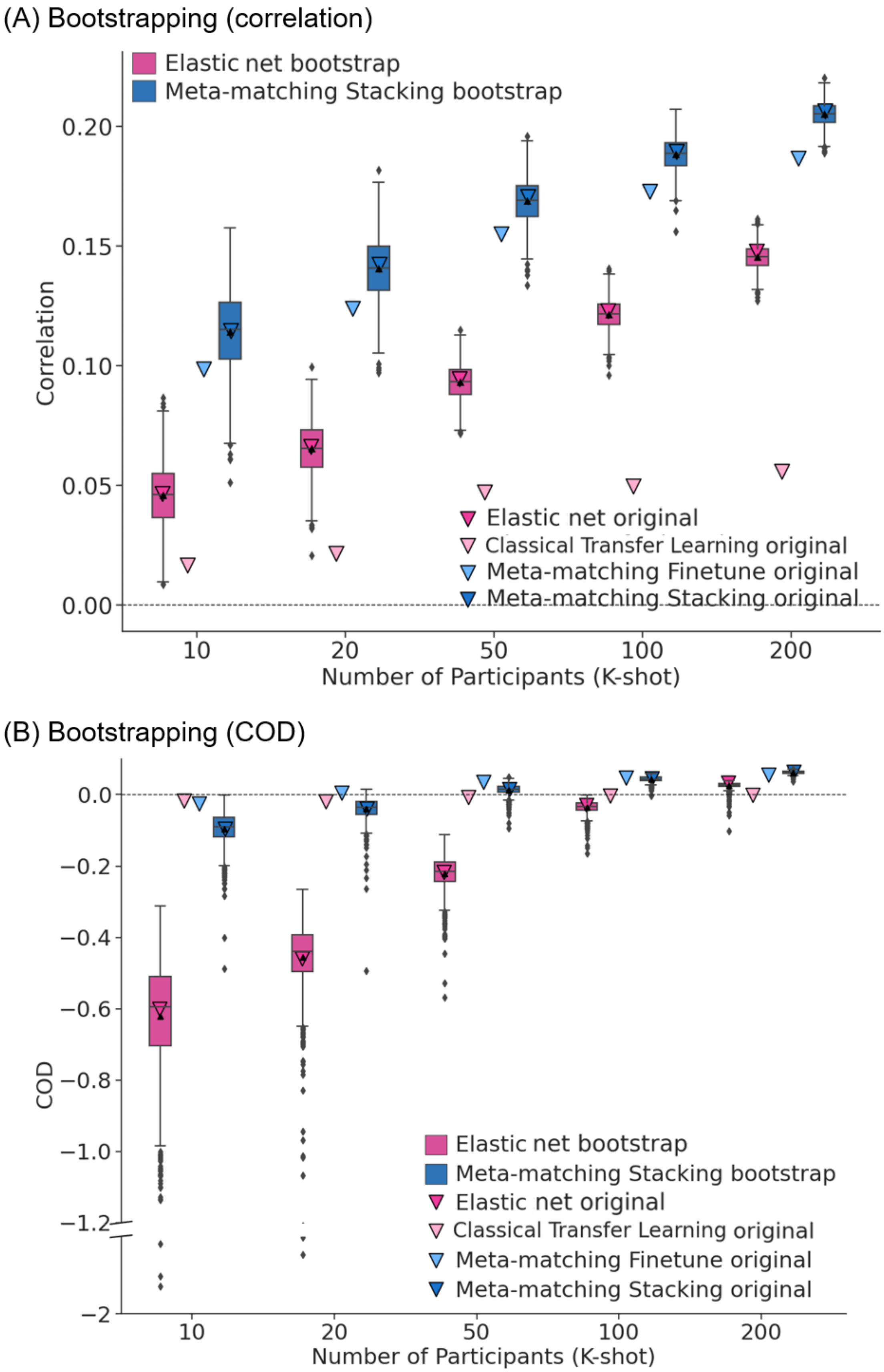

(A) Bootstrapping (correlation)
Elastic net bootstrap
Meta-matching Stacking bootstrap
Elastic net original
Classical Transfer Learning original
Meta-matching Finetune original
Meta-matching Stacking original
Correlation
0.00
0.05
0.10
0.15
0.20
10
20
50
100
200
Number of Participants (K-shot)
(B) Bootstrapping (COD)
Elastic net bootstrap
Meta-matching Stacking bootstrap
Elastic net original
Classical Transfer Learning original
Meta-matching Finetune original
Meta-matching Stacking original
COD
0.0
−0.2
−0.4
−0.6
−0.8
−1.0
−1.2
−2
10
20
50
100
200
Number of Participants (K-shot)

**Figure S1**. Meta-matching outperformed elastic net and classical transfer learning within UK Biobank bootstrapping experiment. (A) Phenotypic prediction performance (Pearson's correlation) (averaged across 34 meta-test phenotypes) on bootstrapped samples in the meta-test set of UK Biobank. X-axis is the number of participants in the meta-test set of UK Biobank used to train an elastic net baseline or adapt the pretrained model from the meta-training set of UK Biobank. Boxplot shows the distribution of performance over 1,000 repetitions of bootstrapped K participants. Inverted triangles represent the average prediction performance (Pearson's correlation) on 100 repetitions of K participants. (B) Same analysis for COD.

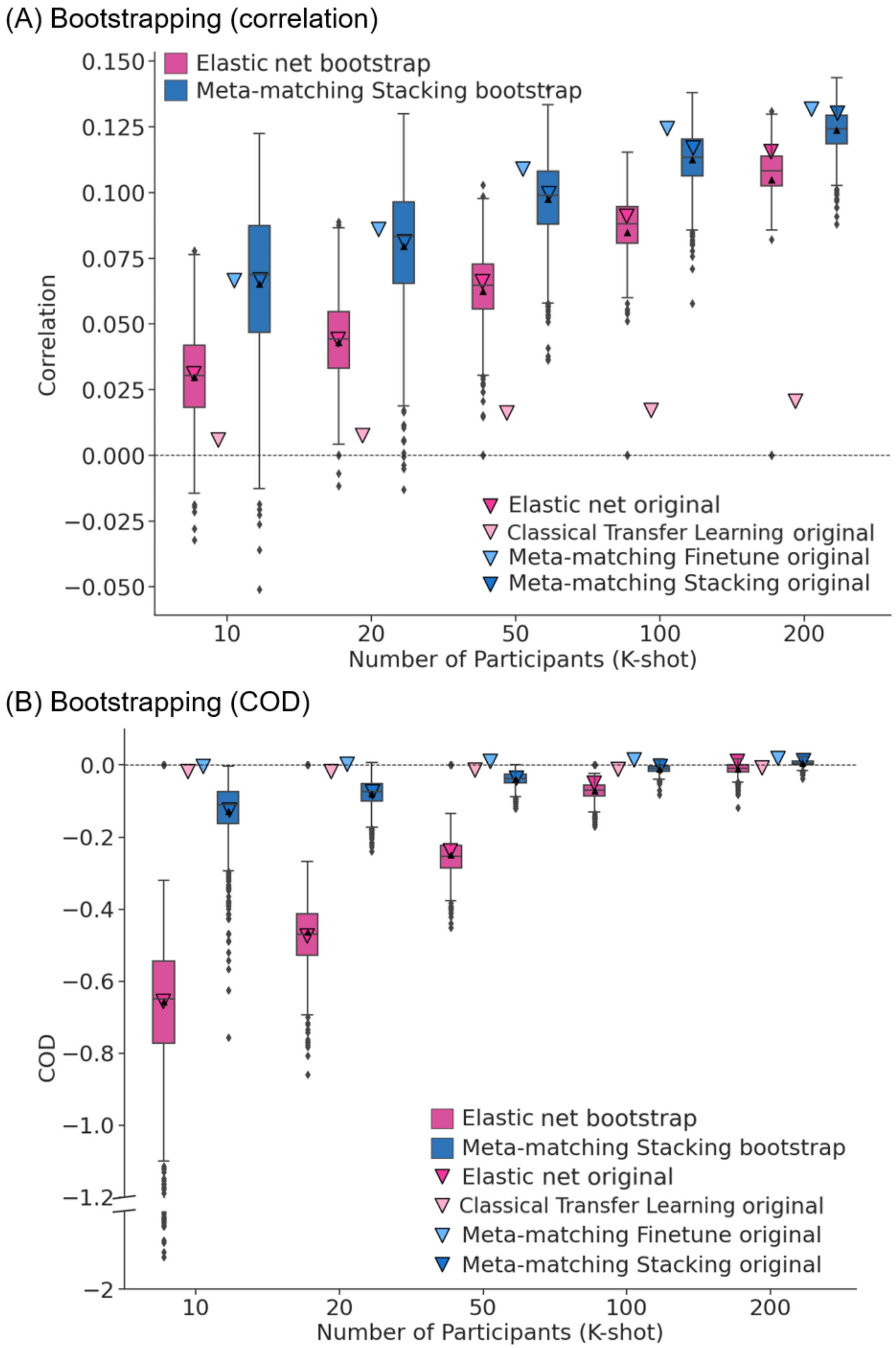

(A) Bootstrapping (correlation)
Elastic net bootstrap
Meta-matching Stacking bootstrap
Correlation
0.150
0.125
0.100
0.075
0.050
0.025
0.000
−0.025
−0.050
Elastic net original
Classical Transfer Learning original
Meta-matching Finetune original
Meta-matching Stacking original
10
20
50
100
200
Number of Participants (K-shot)
(B) Bootstrapping (COD)
COD
0.0
−0.2
−0.4
−0.6
−0.8
−1.0
−1.2
−2
Elastic net bootstrap
Meta-matching Stacking bootstrap
Elastic net original
Classical Transfer Learning original
Meta-matching Finetune original
Meta-matching Stacking original
10
20
50
100
200
Number of Participants (K-shot)

**Figure S2.** Meta-matching outperformed elastic net and classical transfer learning for HCP-YA bootstrapping experiment. (A) Phenotypic prediction performance (Pearson's correlation) (averaged across 35 meta-test phenotypes) on bootstrapped samples in the HCP-YA dataset. X-axis is the number of participants in the HCP-YA dataset used to train an elastic net baseline or adapt the pretrained model from the meta-training source dataset (UK Biobank). Each boxplot shows the distribution of performance over 1,000 repetitions of bootstrapped K participants. (B) Same analysis for COD.

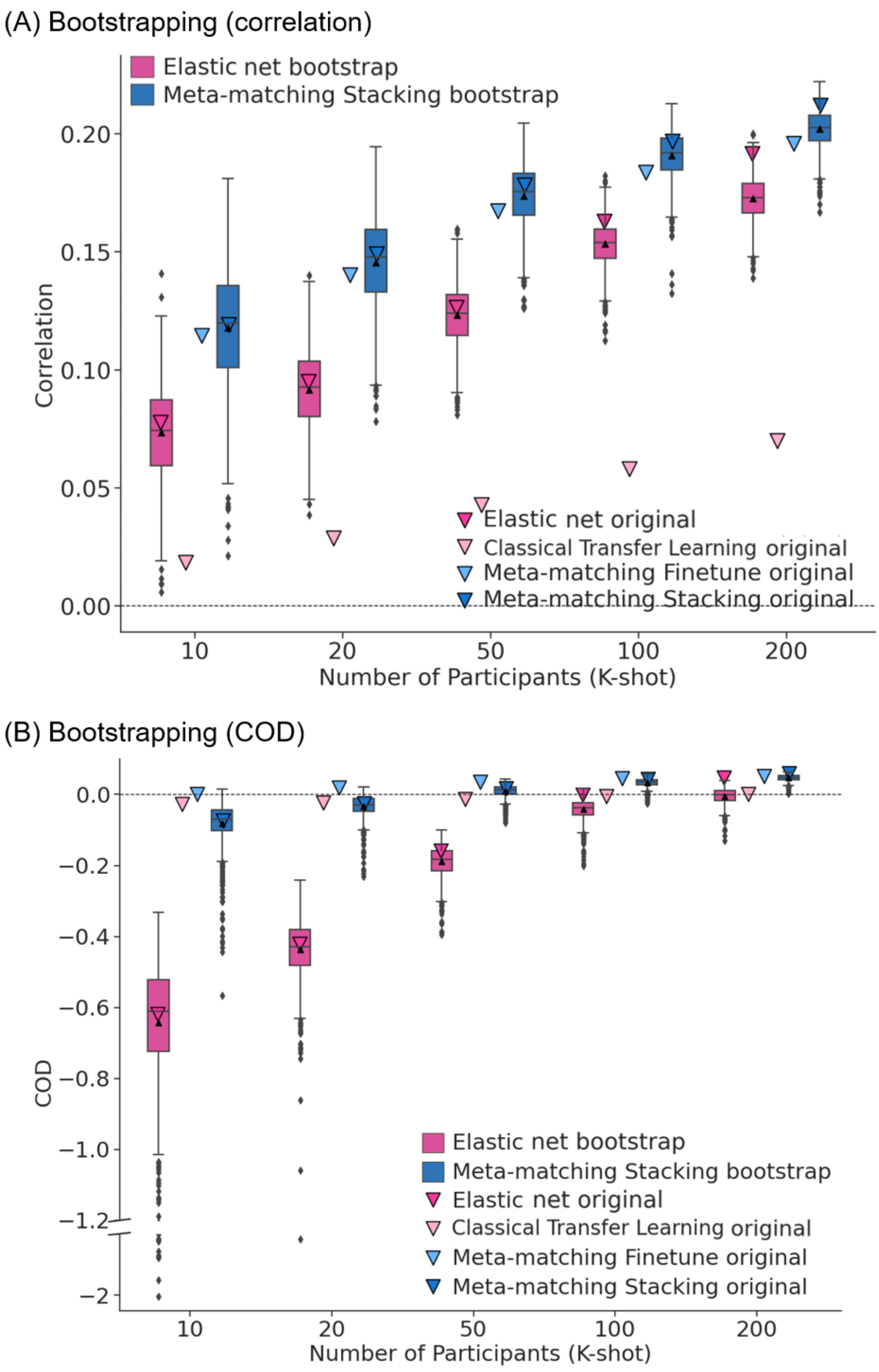
(A) Bootstrapping (correlation)
Elastic net bootstrap
Meta-matching Stacking bootstrap
Elastic net original
Classical Transfer Learning original
Meta-matching Finetune original
Meta-matching Stacking original
Correlation
0.20
0.15
0.10
0.05
0.00
10
20
50
100
200
Number of Participants (K-shot)
(B) Bootstrapping (COD)
COD
0.0
−0.2
−0.4
−0.6
−0.8
−1.0
−1.2
−2
Elastic net bootstrap
Meta-matching Stacking bootstrap
Elastic net original
Classical Transfer Learning original
Meta-matching Finetune original
Meta-matching Stacking original
10
20
50
100
200
Number of Participants (K-shot)

**Figure S3.** Meta-matching outperformed elastic net and classical transfer learning for HCP-Aging bootstrapping experiment. (A) Phenotypic prediction performance (Pearson's correlation) (averaged across 45 meta-test phenotypes) on bootstrapped samples in the HCP-Aging dataset. X-axis is the number of participants in the HCP-Aging dataset used to train an elastic net baseline or adapt the pretrained model from the meta-training source dataset (UK Biobank). Each boxplot shows the distribution of performance over 1,000 repetitions of bootstrapped K participants. (B) Same analysis for COD.
